# Interpretable deep learning reveals the sequence rules of Hippo signaling

**DOI:** 10.1101/2024.02.22.580842

**Authors:** Khyati Dalal, Charles McAnany, Melanie Weilert, Mary Cathleen McKinney, Sabrina Krueger, Julia Zeitlinger

## Abstract

The response to signaling pathways is highly context-specific, and identifying the transcription factors and mechanisms that are responsible is very challenging. Using the Hippo pathway in mouse trophoblast stem cells as a model, we show here that this information is encoded in *cis*-regulatory sequences and can be learned from high-resolution binding data of signaling transcription factors. Using interpretable deep learning, we show that the binding levels of TEAD4 and YAP1 are enhanced in a distance-dependent manner by cell type-specific transcription factors, including TFAP2C. We also discovered that strictly spaced *Tead double* motifs are widespread highly active canonical response elements that mediate cooperativity by promoting labile TEAD4 protein-protein interactions on DNA. These syntax rules and mechanisms apply genome-wide and allow us to predict how small sequence changes alter the activity of enhancers *in vivo*. This illustrates the power of interpretable deep learning to decode canonical and cell type-specific sequence rules of signaling pathways.

**Graphical abstract:** 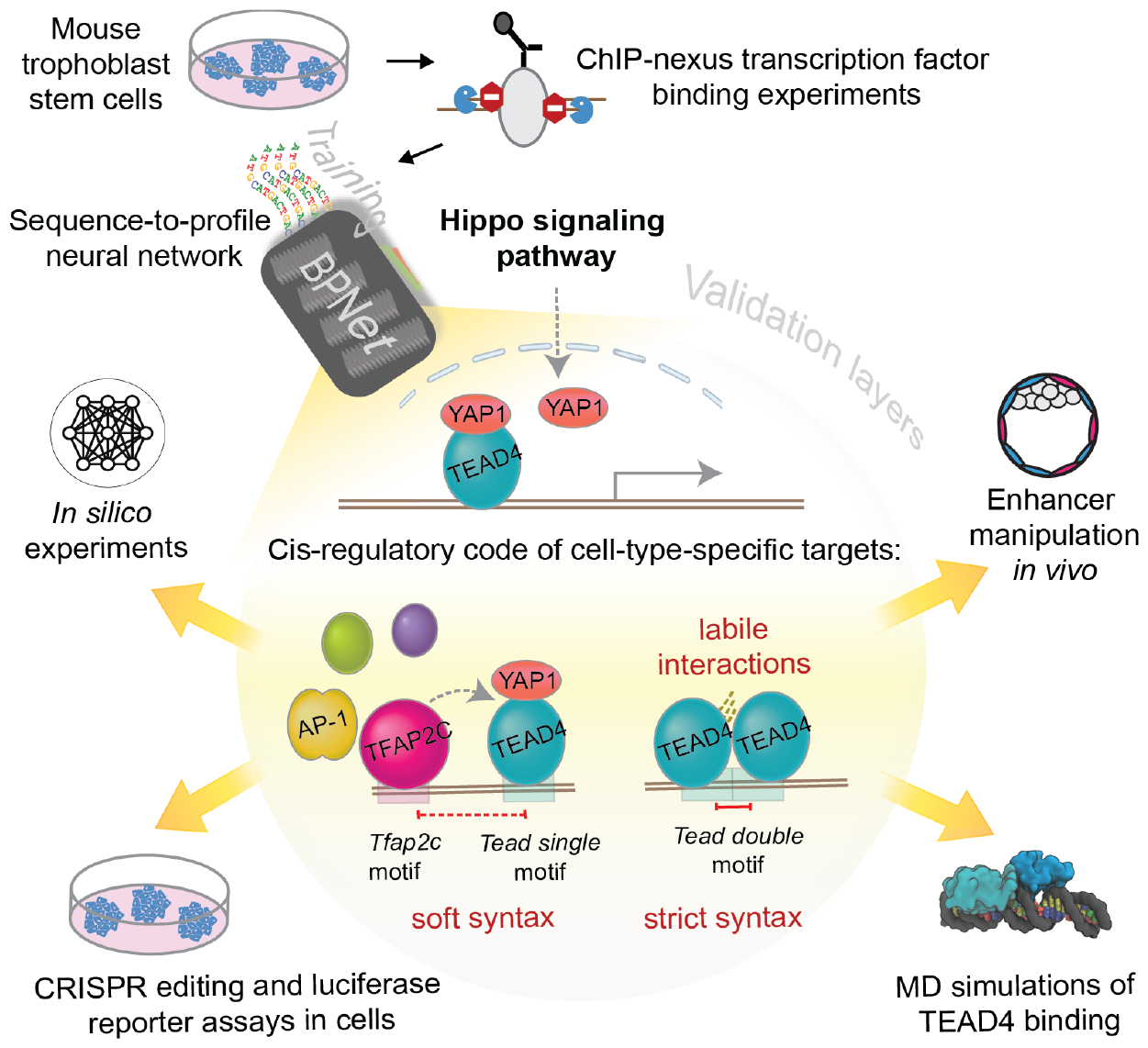

## Introduction

Signaling pathways are critical for cell fate decisions during development, the generation of cell types *in vitro*, and therapeutic interventions, which often target specific signaling pathways^1^. While signal transduction components are typically well studied, how signaling pathways regulate target genes in a cell-type-specific fashion is a fundamental gap in knowledge. Once a signal is transduced into the nucleus, activated transcription factors (TFs) recognize specific DNA sequence motifs or find obligate partner TFs with DNA binding specificity, but which of the *cis*-regulatory sequences become active enhancers and regulate target genes is poorly understood^2,3^. Thus, signaling pathways are critical for gene regulation, but their target specificity is one of the least understood areas of enhancer biology, making it difficult to predict the activity of enhancers or modify their function during development through targeted mutations^3–7^.

A critical question is how signaling TFs interact with other sequence-specific TFs present in that cell type. Signaling pathways are reiteratively used during development, and target genes are regulated depending on cell-type-specific TFs^2,8^. Indeed, cell-type-specific TFs help determine where signaling TFs bind^9–13^, suggesting that signaling pathways rely on TF cooperativity to regulate target genes. However, which TFs can become such partners and which molecular mechanisms are used to help the binding of signaling TFs is poorly understood^14^.

Here, we hypothesized that this TF cooperativity is DNA sequence-driven and thus can be studied by measuring the binding of TFs on DNA and identifying the underlying sequence rules using interpretable deep learning. During training, deep learning models accurately learn sequence rules within genomic regions in an inherently combinatorial manner *de novo* until they can predict the data from sequence alone^15–21^. The key step is then to interrogate the model and extract the learned sequence rules using interpretation tools^15^. This reveals the learned TF motifs, including measurements of their relative affinities and the syntax rules, thus the distance relationships by which motifs cooperate with each other^15,17,20,22,23^.

Interpretable deep learning should, therefore, allow us to identify potential syntax rules by which cell type-specific TFs contribute to the response to signaling while also providing clues on the mechanisms. For example, TF cooperativity that depends on two motifs being spaced at a specific distance is known from the enhanceosome model, where protein-protein interactions stabilize the complex^24–26^. On the other hand, strictly spaced motifs are not frequently observed in the genome, raising the question of whether TF binding cooperativity may also occur through more flexible motif syntax^27–29^. For example, we previously found that a TF may enhance the binding of another TF through soft motif syntax, which occurs at variable motif distances within ∼150 bp but is stronger at closer distances^15,20^.

Whether such syntax rules exist for signaling TFs has previously been difficult to decipher. ChIP-seq binding data tend to be of low resolution and display low levels of signal when the TF binds indirectly through a partner TF^10,11,13^. Likewise, individually manipulating enhancer sequences *in vivo* limits throughput, and the effects can be difficult to interpret since they may be enhancer-specific or caused by the inadvertent disruption of other important sequences^30,31^. Large-scale reporter assays, on the other hand, have produced conflicting results on whether motif syntax is important and have not revealed whether synergistic effects of motifs are mediated through cooperative binding^16,27,32–36^. For these reasons, TF binding cooperativity downstream of signaling pathways has not been systematically studied from a sequence perspective.

To discover these potential sequence rules, we performed the TF binding experiments at the highest resolution and leveraged our previously developed deep learning model BPNet to predict the data at base resolution from genomic sequences of 1-kb^15,20,22,37,38^. This approach optimally resolves sequence rules between closely spaced motifs within enhancers^15^. Since the model does not predict enhancer activity or target genes, we evaluated and validated these downstream aspects using traditional methods.

As a model system, we studied the Hippo signaling pathway in mouse trophoblast stem cells (TSCs). Hippo signaling is critical for specifying trophoblast versus inner cell mass cell fate in the early mouse embryo^39–44^. When cells of the embryo sense that they are facing the outside, i.e., less cell density, they polarize and inactivate the Hippo pathway. This causes YAP1 to translocate to the nucleus and bind to TEAD4, which, like all TEAD family members, binds to a consensus *Tead* motif^40,42,45^. In addition to the Hippo pathway, additional TFs are known to be important for TSC identity, including CDX2, TFAP2C, and GATA3^46–53^, making TSCs an ideal system to dissect the interactions between Hippo signaling TFs and cell type-specific TFs in enhancer activation.

We found that the Hippo pathway effector YAP1 indeed depends on TF cooperativity, whose genome-wide syntax rules can be uncovered by interpretable deep learning. While YAP1 binds directly to TEAD4, the interactions with other TFs are sequence-driven and occur through two distinct mechanisms that manifest themselves through soft or strict motif syntax. We show and experimentally validate that TFAP2C enhances YAP1/TEAD4 binding through distance-dependent soft motif syntax, while two TEAD4 proteins also cooperate through labile protein-protein interactions in the presence of a strictly-spaced *Tead double* motif. This highly cooperative and active motif is surprisingly widespread and has been missed as a canonical element of the Hippo pathway because it is highly variable in sequence. This demonstrates how deep learning models can uncover precise sequence rules by which signaling TFs mediate cell type-specific effects.

## Results

### Deep learning model reveals binding motifs for Hippo TFs

We generated genome-wide, high-resolution binding data for the Hippo signaling TFs (TEAD4 and YAP1) and the potential TSC-specific partner TFs (CDX2, TFAP2C, and GATA3) by using a ChIP-exo technique called ChIP-nexus^37^, in which an exonuclease step generates narrow and sharp binding footprints (Figures 1A, S1A). We used TSCs derived from mouse blastocysts^54^ and confirmed that they retain features of endogenous trophectoderm cells by reintegrating them into the trophectoderm layer of blastocyst embryos in an aggregation assay (Figure S1B). The ChIP-nexus binding data revealed that YAP1 and TEAD4 were more correlated with each other than any other TF pair (Figure 1B), consistent with YAP1 binding to DNA through TEAD4^55–57^.

**Figure 1:**
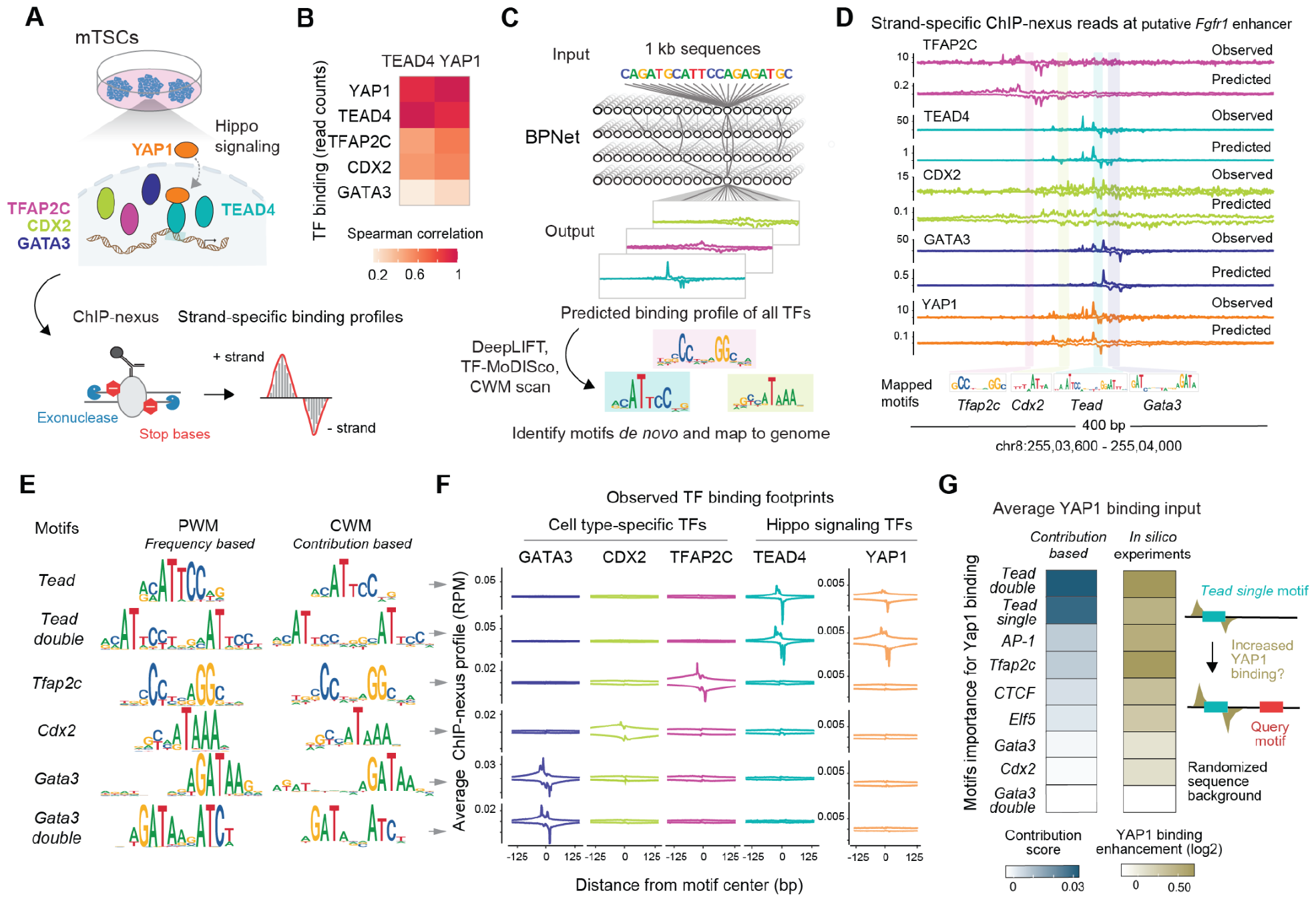
BPNet suggests combinatorial binding motifs for Hippo TFs. **A**) Experimental design to map the high-resolution binding of signaling and cell type-specific TFs in mouse trophoblast stem cells. **B**) Spearman correlations of the ChIP-nexus read counts between TFs at non-promoter binding regions show that YAP1 and TEAD4 binding are highly correlated. **C**) Schematic of the multi-task BPNet model trained to predict ChIP-nexus TF binding footprints based on DNA sequence alone. The interpretation tools DeepLIFT and TF-MoDISco identify and map the motifs for each TF. **D**) The similarity between experimentally observed and BPNet-predicted binding for each TF at the withheld *Fgfr1* enhancer (mm10-chr8:25503600-2550400) illustrates BPNet’s predictive accuracy. Experimental and predicted data each show the + strand on top and the - strand at the bottom. BPNet-mapped motifs for this region are shown below. **E**) Learned motifs are shown as frequency-based position weight matrix (PWM) and contribution weight matrix (CWM), where the base height reflects the contribution to the TF binding predictions. The two motif representations are highly similar for all TFs. **F**) Average ChIP-nexus binding footprints of all TFs at BPNet-mapped motifs. The binding profiles are centered on motifs and shown as read per million (RPM) by positive values on the + strand on top and negative values on the - strand at the bottom. Sharp footprints typically indicate direct binding of the TF to the motif. YAP1 also has sharp footprints on the two *Tead* motifs despite binding indirectly. **G**) Contribution of each motif to YAP1 binding as extracted from the trained BPNet model. Average profile contribution scores as derived by DeepLIFT are shown for each motif in blue (left), while the average binding enhancement of YAP1 as derived from *in silico* experiments in a randomized background are shown in olive (right). Briefly, upon injecting *Tead single* motif (ACATTCCTG) into randomized sequences, other TF motifs were injected at a given distance away for up to 150 bp, and the average predicted YAP1 binding enhancement over no added side query motif was calculated (right plot).

We then trained the deep learning model BPNet to predict the base-resolution binding profiles of all TFs from 187,775 reproducibly bound genomic regions (Figure 1C) by separating chromosomes into training, validation, and test groups to confirm model accuracy^15^. For all TFs, we obtained high prediction accuracy for the read counts, as well as footprint positions on par with the similarity between replicate experiments (Figures S1C-D). We confirmed the results through cross-validation on different chromosome combinations, ensuring model stability (Figure S1E).

After model validation, we extracted the learned motifs^15^. Using an attribution method^58^, we assigned contribution scores to all bases in the input sequences and then summarized *de novo* learned motifs as a contribution weight matrix (CWM)^59^. The CWMs are then used to label motif instances in each genomic region (Figure 1C). Since this mapping approach relies on contribution scores, the mapped motifs depend on the surrounding genomic context learned by the model, thereby outperforming traditional mapping techniques that entirely rely on match scores from a position weight matrix (PWM)^15^.

As a result, the mapped motifs are highly congruent with experimentally derived TF footprints, as illustrated by putative enhancers for *Fgfr1*^42,60^ (Figure 1D), *Amotl2, Pard3b* and *Krt8/18*^61–63^ (Figures S1F-H). These genomic regions were not seen by the model during training, yet the predicted ChIP-nexus profiles are very similar to the experimental data, with footprints found around mapped motifs (Figures 1D, S1F-G). Notably, BPNet also predicted clear ChIP-nexus binding footprints for YAP1, which lacks a DNA binding domain^57,64^(Figures 1D, S1F-H). This is consistent with the potential of neural networks to denoise data^65^.

Among the discovered motifs were the known consensus motifs of the profiled TFs (Figure 1E) and two unexpected motifs: a *Tead double* motif and a *Gata3 double* motif (Figure 1E). These motifs are directly bound by the corresponding TFs, as confirmed by the sharpness of the average ChIP-nexus footprints (Figure 1F). Interestingly, YAP1 also showed sharp binding footprints on both *Tead* motifs in the averaged experimental data. This is consistent with the sharp binding footprint of YAP1 in the predictions and suggests a tight physical association between YAP1 and TEAD4 on DNA. In addition, we identified motifs for TFs that we did not profile, including AP-1 (Jun/Fos), CTCF, and ELF5 (Figure S1E), suggesting that they help the profiled TFs bind.

We next focused on YAP1 binding and analyzed whether and how much, on average, each motif contributes to the YAP1 binding predictions (Figure 1G). The *Tead* motifs, which we will refer to as *Tead single* and *Tead double* motifs, both had a strong contribution, as expected, but the *AP-1* and *Tfap2c* motifs also had a sizable contribution to YAP1 binding (Figures 1G). TFAP2C has a well-established role in trophectoderm specification^46,47,53,66–68^, and AP-1 cooperates with TEAD and YAP in cancer cell lines^69–72^. These data support the model’s learned contribution of the *AP-1* and *Tfap2c* motifs to YAP1 binding and suggest that AP-1 has an earlier role in trophoblast cells than previously implicated^73^.

However, we noted that the model did not assign all motifs the same importance (Figure 1G). For example, CDX2 and GATA3 were profiled because they are critical for trophoblast identity^49–51^, yet neither motif was predicted to help TEAD4 and YAP1 bind. To internally validate these results, we used a second interpretation method, *in silico* synthetic analysis, to systematically test the sequence rules learned by the model (Figure 1G). We injected a *Tead single* motif with or without other motifs into a randomized sequence background and let the model predict how much any given motif enhanced the binding of YAP1 to the *Tead single* motif. The results were similar to the previously extracted contribution scores (Figure 1G, left), which validates our model interpretation and lets us conclude that there are learnable rules by which cell type-specific TFs contribute to the binding of signaling TFs.

### YAP1 binding correlates with markers of enhancer activity

Having analyzed TSC-specific YAP1 binding, we wanted to investigate whether high YAP1 binding levels are indicative of enhancer activation. We expect YAP1 to be a strong activator based on previous molecular evidence^74–76^, but many other TFs have transactivation domains, and thus, it is unclear how much YAP1 contributes to enhancer activation at a genome-wide level.

Since no individual assay unambiguously measures enhancer activity^77^, we performed experiments in TSCs to profile multiple markers of active enhancers: ChIP-nexus for RNA Polymerase II (Pol II), TT-seq to capture enhancer transcription, ATAC-seq to measure chromatin accessibility, and ChIP-seq for H3K27ac found on nucleosomes flanking active cell-specific enhancers with TF-bound motifs^78–80^ (Figure S2A).

Based on these data, active enhancers are indeed associated with particularly high levels of YAP1 binding, as illustrated at an enhancer downstream of the *Bmp7* gene (Figures 2A, S2E-H). TFAP2C, TEAD4, and YAP1 show strong binding footprints, and the contribution scores show that *Tfap2c* and *Tead single* motifs were used by BPNet to predict YAP1 binding (Figure 2A). This region possesses all the characteristic features of active enhancers, with central ATAC-seq accessibility, flanking H3K27ac signal, Pol II occupancy, and bidirectional nascent RNA transcription (Figure 2A).

**Figure 2:**
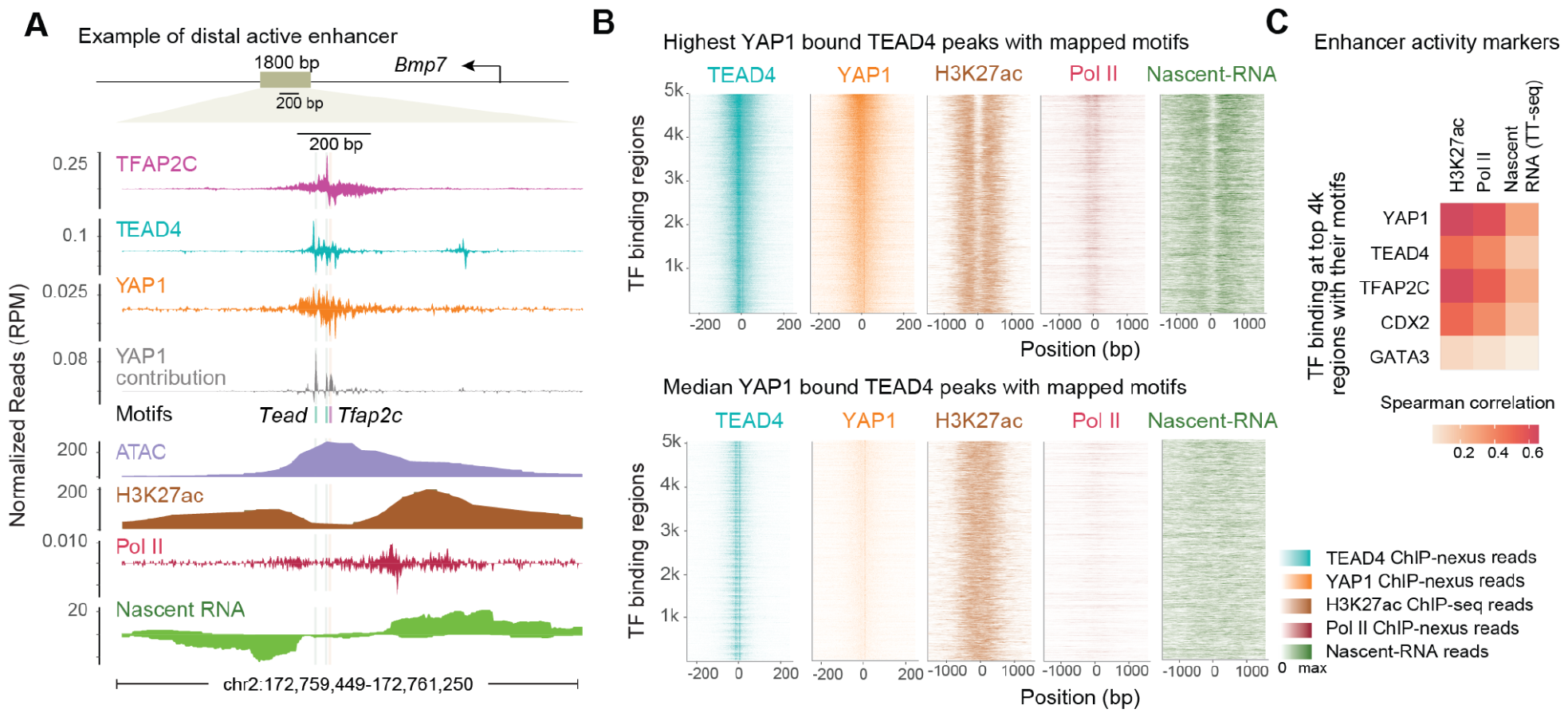
YAP1 binding correlates with enhancer activity markers. **A**) An example of an active enhancer ∼100 kb downstream of the *Bmp7* gene, showing ChIP-nexus TF binding for TFAP2C, TEAD4, and YAP1 alongside BPNet-mapped motifs *Tead* and *Tfap2c* and predicted YAP1 binding contribution. The additional tracks are the fragment coverage for ATAC-seq, H3K27ac ChIP-seq, stranded Pol II ChIP-nexus data, and nascent-RNA-seq derived from TT-seq. **B**) Profile heatmaps of TEAD4 and YAP1 ChIP-nexus data at the 5,000 TEAD4 peaks with the highest YAP1 binding (top) and 5,000 peaks with median YAP1 binding (bottom). Each row represents a region with normalized signal intensity. Note the active enhancer signature of H3K27ac ChIP-seq, Pol II ChIP-nexus, and stranded Nascent-RNA reads at the regions with the highest YAP1 (top). **C**) A heatmap depicting Spearman correlations between ChIP-nexus TF binding and enhancer activity markers (H3K27ac, Pol II, and nascent RNA) at the top 4000 non-promoter peaks containing their corresponding motif. YAP1 correlates best, followed by TFAP2C.

We identified thousands of active enhancers in this way and focused on active enhancers near important trophoblast genes for further characterization. Named after the corresponding gene, these include a *Rin3, Ezr, Cited2, Amotl2, Bmp7, Dst*, and *Tjp1* enhancer, respectively. They were validated by cloning the minimal central region into a luciferase reporter assay and measuring their activity in TSCs (Figure S2B).

To visualize the global correlation between YAP1 binding and the markers of enhancer activity, we selected all TEAD4 peaks containing its mapped motifs; we then compared the 5,000 regions with the highest YAP1 binding to 5,000 regions with median levels of YAP1 binding (Figure 2B). The top YAP1 bound regions showed strong H3K27ac signal, Pol II binding, and nascent transcription adjacent to the central region, while no strong evidence of enhancer activity was observed for the more lowly bound set (Figure 2B).

Finally, we directly examined the correlation between each TF’s binding and enhancer activity markers (H3K27ac, Pol II, and nascent RNA) (Figure 2C). Using each TF’s top 4,000 non-promoter ChIP-nexus peaks containing their motifs, we found that among all TFs, YAP1 binding correlated best with the enhancer activity markers. Given the strong transactivation potential of YAP1^74–76,81^, we conclude that YAP1 binding is an important determinant for enhancer activation in TSCs.

### Enhancer activation involves DNA distance-dependent TF cooperativity

If the binding of YAP1 can both determine enhancer activation and be enhanced by cell-specific TFs, we would expect the corresponding motifs of cell-specific TFs to activate transcription synergistically. Synergistic activation by two motifs has been documented^16,82–84^, but the mechanisms are not clear and could vary. Hence, it is important to understand whether YAP1’s cooperativity with cell type-specific TFs is mechanistically connected to its ability to drive enhancer activation. To address this, we experimentally tested whether the BPNet-derived syntax rules regulate enhancer activity.

We focused on *Tead single* (to which YAP1 binds through TEAD4) and *Tfap2c* motifs since this motif pair is frequently found at active enhancers (Figure 2). Additionally, genes near these active enhancers are enriched for cell fate commitment and GTPase regulation (Figure S2C), consistent with previous studies^42,50,53,66^.

We first tested for synergistic activation by performing luciferase assays using the 200 bp minimal *Bmp7* enhancer, which has a *Tead single* and *Tfap2c* motif (shown in Figure 2A). We perturbed combinations of these motifs by mutating the two bases of each motif that contributed most to the predictions and confirming through our model that this led to decreased YAP1 binding (Figure S2D). Luciferase assays showed that mutating either motif alone was sufficient to strongly reduce the activity while mutating both almost completely abolished the activity (Figure 3A). Thus, each motif in the *Bmp7* enhancer produced only moderate activity alone, while together, they resulted in activity that exceeded the sum of each motif’s effect. These results show that the *Tead single* and *Tfap2c* motifs mediate activation synergistically, presumably at least in part by increasing YAP1 binding.

**Figure 3:**
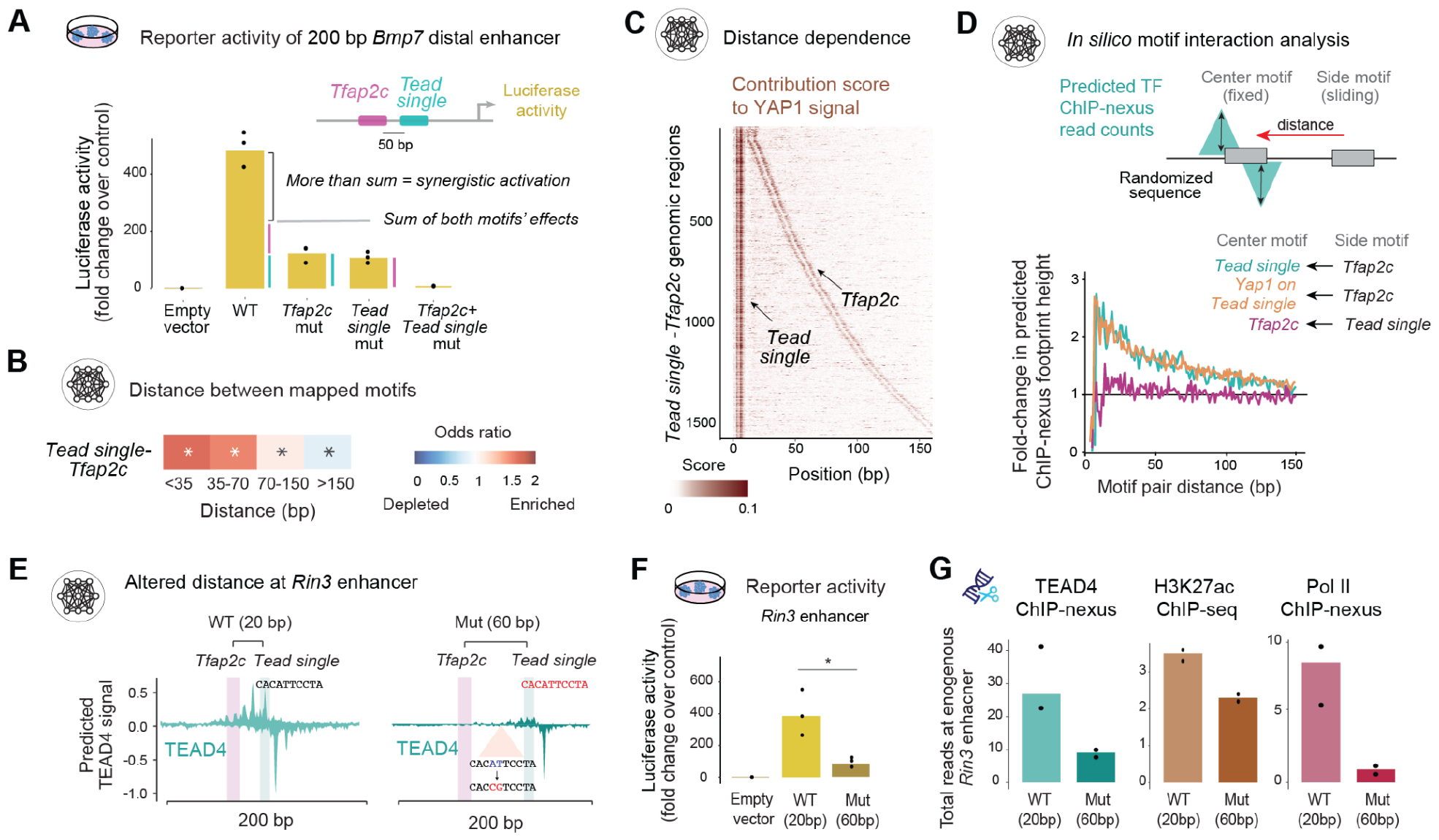
Enhancer activation involves DNA distance-dependent cooperativity. **A**) Luciferase assay, performed across three biological replicates, using 200 bp of the *Bmp7* enhancer (mm10*-chr2:172,760,183-172,760,382)* shows that the *Tead single* and *Tfap2c* motifs function synergistically, producing reporter activity greater than the sum each individual motif **B**) *Tead single* and *Tfap2c* motifs important for YAP1 binding are enriched at short distances, calculated as the odds ratio of the frequencies observed for wild-type over permuted regions. Significance was denoted by \**p* < 10^−5^ using Pearson’s chi-squared test. **C**) Heatmap showing BPNet contribution scores of YAP1 binding across regions with one *Tead single* and one *Tfap2c* motif, ordered by the distance between the motifs (up to 160 bp). The contribution from the *Tfap2c* and *Tead single* motif decreases with larger distances. **D**) During *in silico* analysis, motifs are injected into randomized sequences, and BPNet is used to predict the average enhancement of TF binding to its motif (center) in the presence of a side motif ^15^. The results show a strong distance-dependent enhancement of TEAD4 and YAP1 binding in the presence of a *Tfap2c* motif. **E**) Predicted TEAD4 binding at the wild-type *Rin3* enhancer where the *Tfap2c* and *Tead single* motifs are 20 bp apart (left) and after the distance was increased to 60 bp between motifs (right). The motif was moved by inserting a new motif further away and mutating the most important bases within the *Tead single* motif at its original position. **F**) Luciferase assays of the wild type and mutated 200 bp minimal *Rin3* enhancer were performed in three biological replicates and normalized to the empty vector control. Significance was determined by a student’s t-test (*p* < 0.05). **G**) After mutating the endogenous *Rin3* enhancer in the same way through sequential CRISPR, TEAD4 ChIP-nexus binding (left), H3K27ac ChIP-seq levels (center), and Pol II ChIP-nexus occupancy (right) were all reduced compared to wild-type experiments (WT). Measurements are normalized reads per million (RPM) for ChIP-nexus, mean of log2(H3K27ac/WCE reads) in a 2 kb window for ChIP-seq data.

To understand the syntax rules by which *Tead single* and *Tfap2c* motifs recruit YAP1, we tested for soft motif syntax by determining whether shorter distances between the motif pair have a bigger impact on YAP1 binding than longer distances. We found that short distances of <70 bp were strongly overrepresented over longer distances in our genomic regions (Figure 3B, S3A). Moreover, when we examined genomic regions with spacings up to 160 bp, we found that closer distances showed visually stronger contribution scores toward YAP1 binding (Figures 3C, S3C-D). This confirms that TFAP2C enhances YAP1 binding in a distance-dependent manner.

We next investigated whether TFAP2C directly helps the recruitment of YAP1 or whether the effect on YAP1 binding is mediated through binding cooperativity between TEAD4 and TFAP2C (Figure 3D). To distinguish between these possibilities, we performed *in silico* experiments in randomized sequences, where we injected *Tead single* and *Tfap2c* motifs at distances up to 150 bp and recorded BPNet’s predicted binding enhancement of TEAD4, YAP1, or TFAP2C due to the nearby motif. This revealed that YAP1 and TEAD4 binding both depend on the distance of the nearby *Tfap2c* motif, causing both an over 2.5-fold increase in binding when the *Tfap2c* motif is close (Figures 3D, S3B). Notably, the reverse was not necessarily true: TFAP2C binding was not substantially increased (<1.5-fold) in the presence of a nearby *Tead single* motif (Figure 3D) but showed some increase in the presence of a *Tead double* motif (Figures S3B, S3D). The distance dependence and directionality by which the *Tfap2c* motif enhances YAP1 binding is characteristic of soft motif syntax^15^.

To validate the *Tead single-Tfap2c* soft motif syntax, we performed luciferase reporter experiments on the *Rin3, Dst*, and *Adcy7* enhancers. Using BPNet predictions as a guide for designing experiments, we changed the distances between the *Tead single* and *Tfap2c* motifs by deleting a motif through minimal mutations and introducing a new motif at a different location. In all three cases, the reporter activity of the enhancer changed in the expected direction. For example, when we moved the *Tead single* motif in the minimal *Rin3* enhancer further away from the *Tfap2c* motif (from 20 bp away to 60 bp away), BPNet predicted lower TEAD4 binding (Figure 3E). This lower binding mirrored the lower activity measured in the luciferase assay (Figure 3F). Moving the two motifs closer to each other increased the luciferase reporter activity of the minimal *Dst* and *Adcy7* enhancers (Figure S3E).

To confirm that these distance effects are also observed in the genomic context, we performed CRISPR-Cas9-induced mutations using homologous recombination on the endogenous *Rin3* enhancer (Figure S3F). ChIP experiments on this edited cell line confirmed the reduced TEAD4 binding, H3K27ac, and Pol II levels at the *Rin3* enhancer (Figure 3G), while other enhancers remained unchanged (Figure S3G). This demonstrates that changing the distance between motifs through controlled minimal mutations measurably affects enhancer activation in an *in vivo* endogenous context.

### The *Tead double* motif is a canonical element of the Hippo pathway

Having observed TF binding cooperativity with soft syntax, we next examined whether BPNet also learned examples of strict syntax. Indeed, the double motifs for TEAD4 and GATA3 (Figure 1E) could be strictly spaced composite motifs, where two protein domains bind cooperatively through protein-protein interactions^24,85,86^. In support of this, a different palindromic GATA motif is bound by two GATA zinc fingers^87,88^. However, we were particularly interested in the *Tead double* motif and its role in the Hippo signaling pathway.

Close inspection of the literature revealed an interesting conundrum. The *Tead double* motif has typically not been identified when using traditional analysis approaches on genomics data^42,89^ and is not considered a canonical response element of the Hippo pathway^42,90,91^. However, it has been discovered multiple times in the past^76,92–95^. The first characterization occurred on the SV40 enhancer, but its identity remained unclear since it did not resemble the *Tead single* motif^96–100^. Even when the *Tead double* motif was later independently discovered in *Drosophila* and cancer cells^76,92–94^, it was unknown how widespread its role might be in mediating the response to Hippo signaling.

In contrast to previous studies, which identified the motif only rarely and in low numbers, we discovered and mapped thousands of *Tead double* motifs in TSCs. These are not spurious motifs since they have sharp TEAD4 ChIP-nexus footprints and thus are bound *in vivo* (Figure 4A). This suggests that BPNet is uniquely suited to robustly discover such motifs. Notably, BPNet learns motifs through its contribution to TF binding rather than relying on statistical overrepresentation^15^. Indeed, the optimal spacing of the *Tead double* motif is not found more frequently than other spacings in the genome, e.g., a relative spacing of +2 bp is more frequent (Figure 4B). Moreover, BPNet has been shown to learn relative motif affinities^17,20,23,101^ and thus might be adept at learning degenerate versions of a motif.

**Figure 4:**
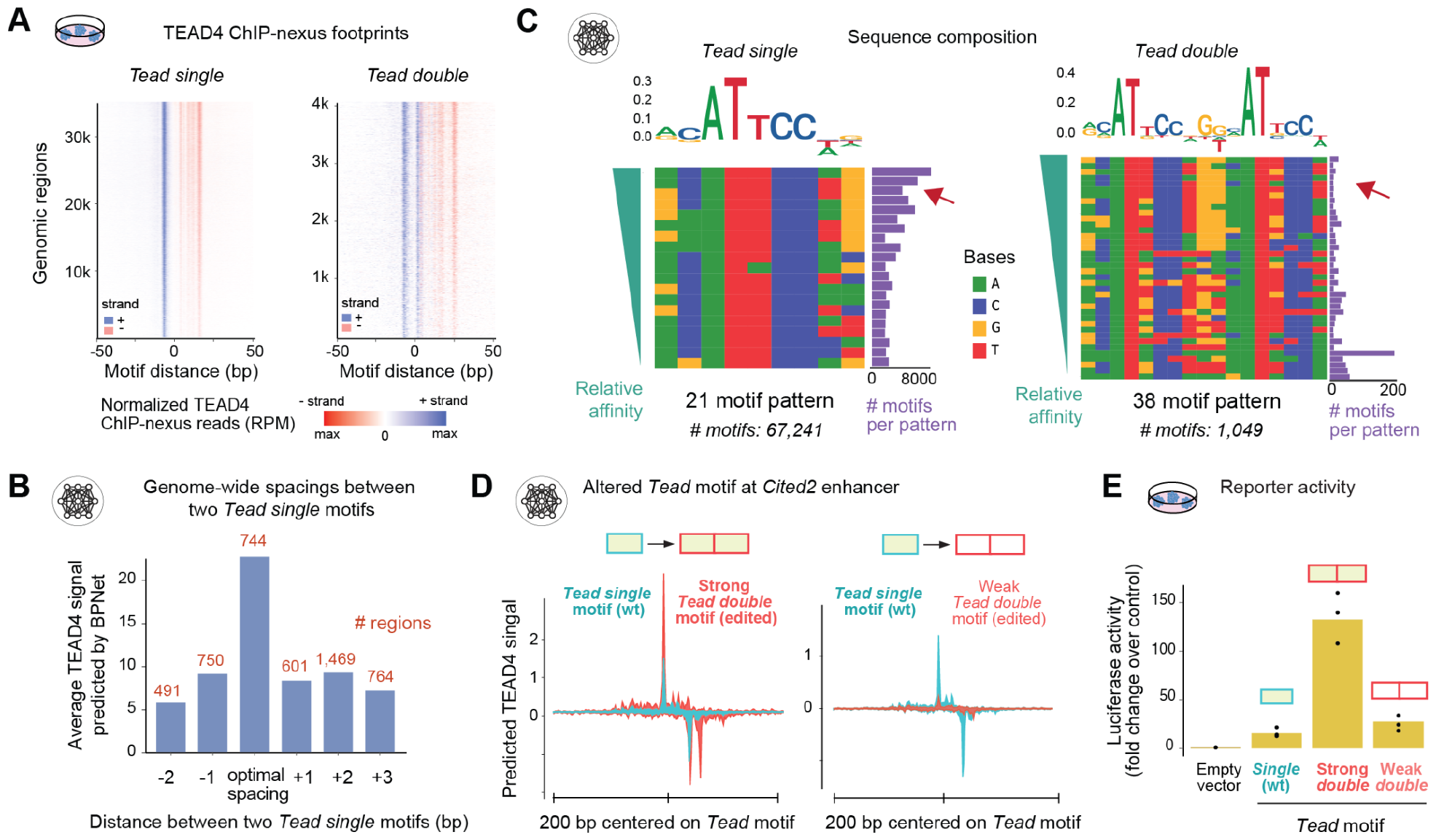
The *Tead double* motif is widespread, highly variable, and active. **A**) BPNet-mapped *Tead single* and *double* motifs are bound by TEAD4 in TSCs *in vivo*, as sharp ChIP–nexus footprints are consistently observed on the + strand (blue) and the - strand (red) for both motifs on regions with ChIP–nexus reads. Distance in bp is shown relative to the left side of the motifs. **B**) BPNet predicts higher TEAD4 binding when two high-affinity *Tead single* motifs have the optimal spacing compared to other spacer lengths, although the optimal spacing is not more frequent (number # of regions shown in red). Predictions were performed after injecting the different motif spaced sequences into random sequences and summed signal in a 50 bp window of the injected motif and averaged across all random sequences. **C**) The *Tead single* and *double* motifs were grouped into sequence patterns shown, where pattern frequency was in the top 90th percentile and equal to or more than 10. The contribution weighted matrix (CWM) logo of the same sequences is shown on top for clarity. Regions were sorted by relative motif affinities (on the left in green), and their frequency is shown (on the right in purple). High-affinity *Tead single* motifs occur most frequently, while *Tead double* motifs do not (red arrow). **D**) At the *Cited2* enhancer, a *single Tead* motif was replaced with a *Tead double* motif, either a strong or a weak one, and the TEAD4 binding profile was predicted by BPNet (200 bp centered on the left side of the motif is shown). **E**) Luciferase assay of the 200 bp minimal *Cited2* enhancer (mm10-chr10:17,579,590-17,579,789) normalized over the empty vector control. Experiments were performed with three different motifs, each with three biological replicates.

Consistent with degeneracy, we found that our mapped *Tead double* motifs had an unusually large number of mismatches to the consensus (Figure 4C). Among the mapped *Tead single* motifs (67,241), the vast majority fall into 21 commonly occurring patterns, with high-affinity motifs being the most frequent (Figure 4C left). In contrast, only 1,049 (6%) of the mapped *Tead double* motifs fall into 38 sequence patterns with 10 or more instances. Thus, the exact sequence pattern of the *Tead double* motif is highly variable (Figure 4C right). Furthermore, the sequence patterns with the highest predicted affinity are not the most frequent (Figure 4C). The highly variable pattern of the *Tead double* motif explains why degenerate versions, e.g., the one on the SV40 enhancer (Figure S4A), are difficult to identify using traditional motif discovery methods, while BPNet learns that they are widespread despite their variability.

Having confirmed the surprisingly widespread occurrence of the *Tead double* motif, we next tested whether the motif mediates enhancer activation (Figures 4D-E). Using the *Cited2* enhancer, we replaced its high-affinity *Tead single* motif with a *Tead double* motif of either higher or lower affinity. BPNet predicted that the strong *Tead double* motif caused a large increase in TEAD4 binding, while the weaker one caused a reduction in binding (Figure 4D). When assayed in a luciferase assay, the strong *Tead double* motif caused an over 8-fold increase in activity compared to the wild-type *Tead single* motif. Interestingly, even the weak *Tead double* motif with low TEAD4 binding showed increased activity (∼1.9 fold) over the *Tead single* motif (Figure 4E). This shows that the *Tead double* motif is active even at lower affinity.

Finally, we asked whether the *Tead double* motif is specific for TSCs or whether it is also bound by TEAD family members in other cell types. When we analyzed BPNet models trained on TEAD1-4 ChIP-seq data from the ENCODE portal^102,103^ (https://www.encodeproject.org/)), we found that BPNet discovered the *Tead double* motif in diverse human cell types (Figure S4B). This suggests that the *Tead double* motif has often been missed because it is variable but that it is, in fact, a widespread canonical motif of the Hippo pathway.

### TEAD4 cooperativity through labile protein-protein interactions

The high binding and activity of the *Tead double* motif likely stems from two TEAD4 molecules binding cooperatively, consistent with previous gel shift assays^92–94,97^. In the simplest scenario, the binding is cooperative because the ternary complex is stabilized on DNA through protein-protein interactions, as observed in crystal structures of other TFs interacting on DNA^104–110^. However, the exact nature of this cooperativity and how it manifests in the genome *in vivo* is not well understood. We therefore, explored the genome-wide rules of cooperativity learned by BPNet while simultaneously conducting all-atom molecular dynamics (MD) simulations to characterize this type of binding cooperativity from a structural perspective (Figure 5).

**Figure 5.**
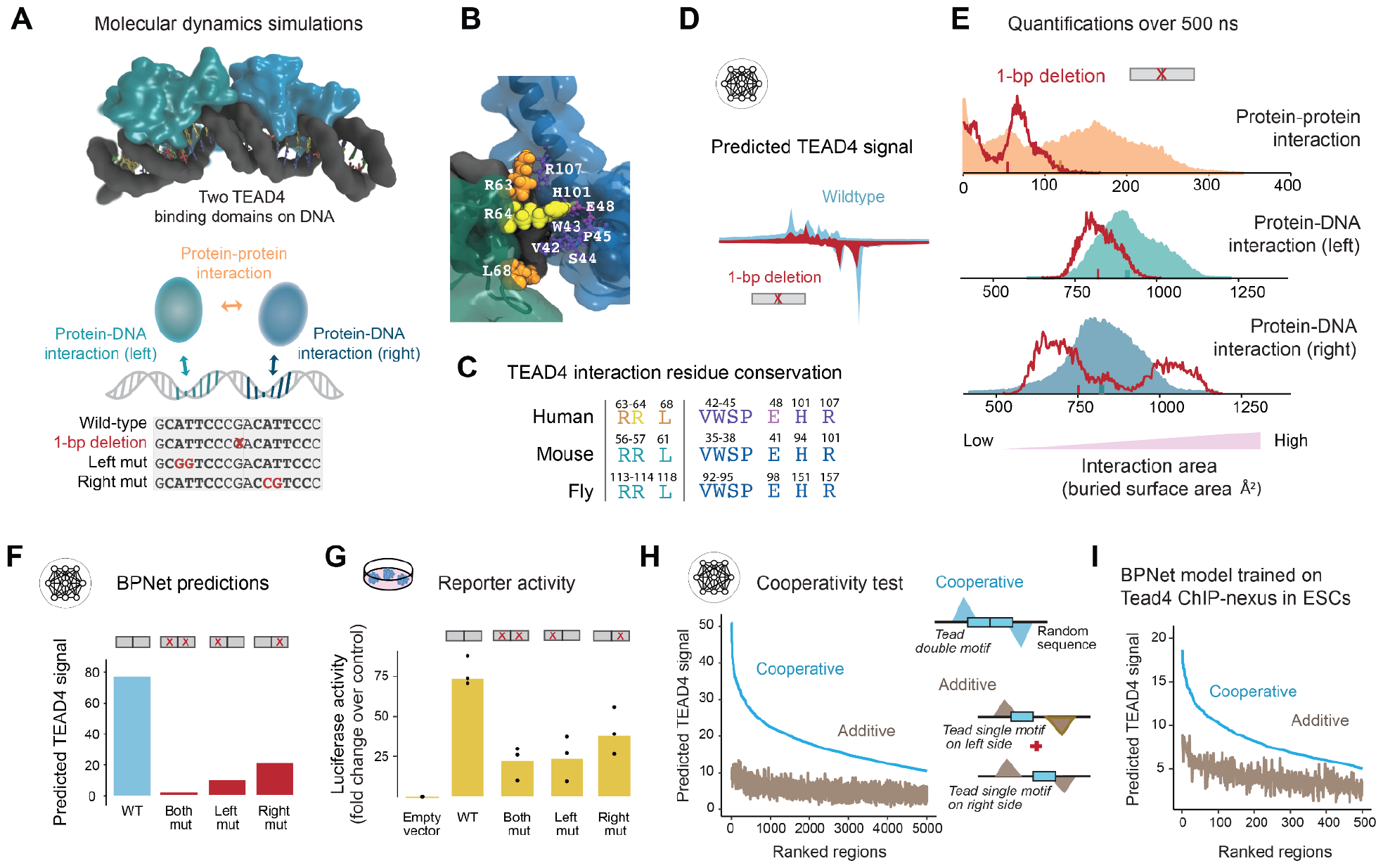
BPNet and MD simulations reveal quantitative details of TEAD4 cooperative binding at double motifs. **A**) Using the known structure of human TEAD4 bound to a single motif^111^, we constructed a model of two TEAD4 DNA binding domains simultaneously bound to a high-affinity double motif (top). The graphic (below) shows which interactions were measured to quantify binding strength. A video of the simulation, showing the dynamics of the protein-protein interaction, is included as supplemental file 1. **B**) Residues involved in interprotein interactions in the TEAD4 DNA binding domain over the course of the simulation. A representative frame magnified shows hydrogen bonding between R64 and E48. Residues closer than 4 Å in at least 20% of simulation frames are shown. **C)** The residues involved in protein-protein interactions are conserved across orthologs of *Drosophila* Scalloped, mouse TEAD4, and human TEAD4, as measured by multiple sequence alignment of Clustal Omega using UniProt id (P30052, Q62296, Q15561). **D**) When injected into random sequences, BPNet predicts lower TEAD4 footprints across the 1-bp deletion, suggesting optimal spacing is important for the cooperativity as in the MD simulations on the right. **E**) Buried surface area distributions from two MD simulations of TEAD4 models. The values from the canonical, high-affinity double motif (GCATTCCCGACATTCCC) are shown as solid areas, and the corresponding values from a 1-bp deletion (GCATTCCCxACATTCCC) are shown as a red line. The protein-protein interaction values are lower when simulating the 1-bp deletion of the TEAD4 double motif, suggesting lower cooperativity between TEAD4 proteins at suboptimal spacing. The mean protein-DNA interaction also decreases in the deletion, suggesting that the whole complex is less stably bound to the DNA. **F**) BPNet predicted TEAD4 binding on the wild-type double motif (GCATTCCCGACATTCCC) and on the same sequence where either the left (GC**GG**TCCCGACATTCCC), the right (GCATTCCCGAC**CG**TCCC), or both (GC**GG**TCCCGAC**CG**TCCC) *Tead single* motif component of *Tead double* are mutated. Predictions were performed after injecting the motif sequences into random sequences and summed signals in a 100 bp window of the injected motif and averaged across all random sequences. **G**) Normalized luciferase assay of the 200 bp genomic region (mm10-*chr7:65,430,487-65,430,686*) consisting of the motif or its mutant variants, as described in D), were performed as three biological replicates. **H**) All CWM-mapped *Tead double* motifs grouped by sequence were injected into random sequences either as a whole or each half to predict the sum signal in a 50 bp window, averaged across all random sequences, and ordered TEAD4 binding in mouse TSCs. **I**) same as in Figure H), but derived *Tead double* motifs and predictions were in mouse ESCs.

The MD simulations were performed by placing two TEAD4 binding domains (PDB:5GZB)^111^ on DNA containing the high-affinity *Tead double* motif from the *Tjp1* enhancer (Figure 5A). This ternary complex remained stable during the simulations over 500 ns, allowing us to quantify the protein-DNA and protein-protein interactions over time (measured as buried surface area Å^2^) (Figure 5B) and identify the interacting amino acid residues, which we found to be an integral part of the TEA DNA-binding domain and highly conserved between TEAD family members and across evolution (Figure 5C). As a control, we performed MD simulations after mutating the *Tead* motif on either side and after changing the spacing between the two *Tead* motifs through a 1-bp deletion or 1-bp addition in the middle (Figures 5A and S4D-F), red line histogram) since BPNet predicted that the TEAD4 binding footprint was strongest on the correctly spaced *Tead double* motif (Figure 5D).

We found that in the presence of the correctly spaced high-affinity *Tead double* motif, the two TEAD4 molecules showed the strongest protein-protein interactions between (Figures 5E, S4D-F). Notably, these interactions changed dynamically on the scale of ∼100 nanoseconds (Video: supplemental file 1) and varied in their molecular details over time (Figure 5E). For comparison, 100 nanoseconds are much faster than the 10s of microseconds seen for stable protein-protein complexes^112^. Furthermore, the measured interaction strength is lower than that between the c-Jun-ATF-2 or RelA-p50 dimers in the enhanceosome crystal structure but similar to the other protein-protein interactions in that structure (Figure S4C). This suggests that the *Tead double* motif serves as a DNA template that facilitates labile interactions between the two TEAD4 binding domains on DNA^24,113^.

To probe for further similarities between our structural and deep learning models, we derived relative motif affinities from each model. This investigation was prompted by an unexpected observation in the MD simulations: although we used a high-affinity *Tead double* motif with identical core *Tead* motifs on each side, we found that in the given orientation, the left motif had stronger interactions with TEAD4 than the right motif (turquoise versus blue histogram, Figure 5E). Therefore, we tested whether BPNet had learned a similar binding asymmetry for the same sequence. When we mutated each *Tead* motif of the *Tead double* motif, BPNet predicted higher TEAD4 binding on the left side than the right (Figures 5F, S4G), consistent with the MD simulations (Figure S4D). This asymmetry was experimentally validated using luciferase assays in the context of the *Tjp1* enhancer (Figure 5G). As an additional example, we analyzed the low-affinity *Tead double* motif in the context of the *Amotl2* enhancer (Figures S4E, S4H-I), which showed a similar asymmetry that we then validated by luciferase assays (Figure S4J).

We next leveraged BPNet’s ability to accurately predict TEAD4 binding on individual *Tead* motifs versus the entire *Tead double* motif to analyze the cooperative binding behavior genome-wide. We did not observe a consistent asymmetry between the two sides of the double motif, indicating that the contribution strength from each side varies. However, we found that *Tead double* motifs showed consistently strong cooperativity, with an average of ∼4-fold higher TEAD4 binding than expected from an additive model of two single *Tead* motifs (Figure 5H).

To test whether the cooperativity was general or specific to TSCs where the Hippo pathway is active, we performed TEAD4 ChIP-nexus experiments in mouse embryonic stem cells (ESCs) where YAP1 is not nuclear^40^. We then trained an independent BPNet model as before but only on TEAD4 ChIP-nexus data in ESCs. We found that TEAD4 binding was lower with fewer bound instances of the *Tead double* motif, but we still observed cooperativity with a >2-fold increase over the additive signal (Figures 5I, S4K). This confirms that some of the cooperativity on the *Tead double* motif is mediated through TEAD4 without requiring YAP1, consistent with the MD model.

In summary, despite being derived independently by very different methods, the MD and BPNet models revealed similar rules on the cooperative binding of TEAD4 on the *Tead double* motif. Both revealed similar binding strengths for the individual *Tead* motifs, showing the strongest cooperativity when combined at optimal spacing in the *Tead double* motif. This is striking, as MD modeling explains TEAD4 cooperativity via labile protein-protein interactions, while the BPNet model exhibits this same cooperativity through predictions of TEAD4 binding across thousands of *Tead double* motifs genome-wide.

### A redesigned enhancer shows that the *Tead double* motif increases gene activation in mouse embryos

As a final validation, we confirmed that the *Tead double* motif mediates the response to the Hippo pathway *in vivo*. Since the Hippo pathway is inactive and YAP1 is nuclear in cells outside the embryo, we added the *Tead double* motif to an endogenous enhancer in TSCs and reintegrated these cells into mouse embryos. We selected the *Ezr* enhancer because it contains a *Tead single* motif, has all the hallmarks of being an active TSC enhancer (Figure S2F), and its target gene *Ezr* encodes an actin-associated protein that is highly expressed in TSCs^53,114,115^. We hypothesized that replacing the *Tead single* motif with a high-affinity *Tead double* motif would increase the enhancer’s activity *in vivo*.

While it is typically easy to destroy the activity of an enhancer by mutating relevant TF motifs^34^, it is more challenging to engineer mutations that increase the activity since sequence changes can have unexpected side effects^30,31^. However, deep learning models are ideal for exploring and evaluating possible mutations, as shown in *Drosophila*^16,116^. We, therefore, used BPNet to replace the *Tead single* motif with a *Tead double* motif while maximizing TEAD4 binding (Figure 6A, left).

**Figure 6:**
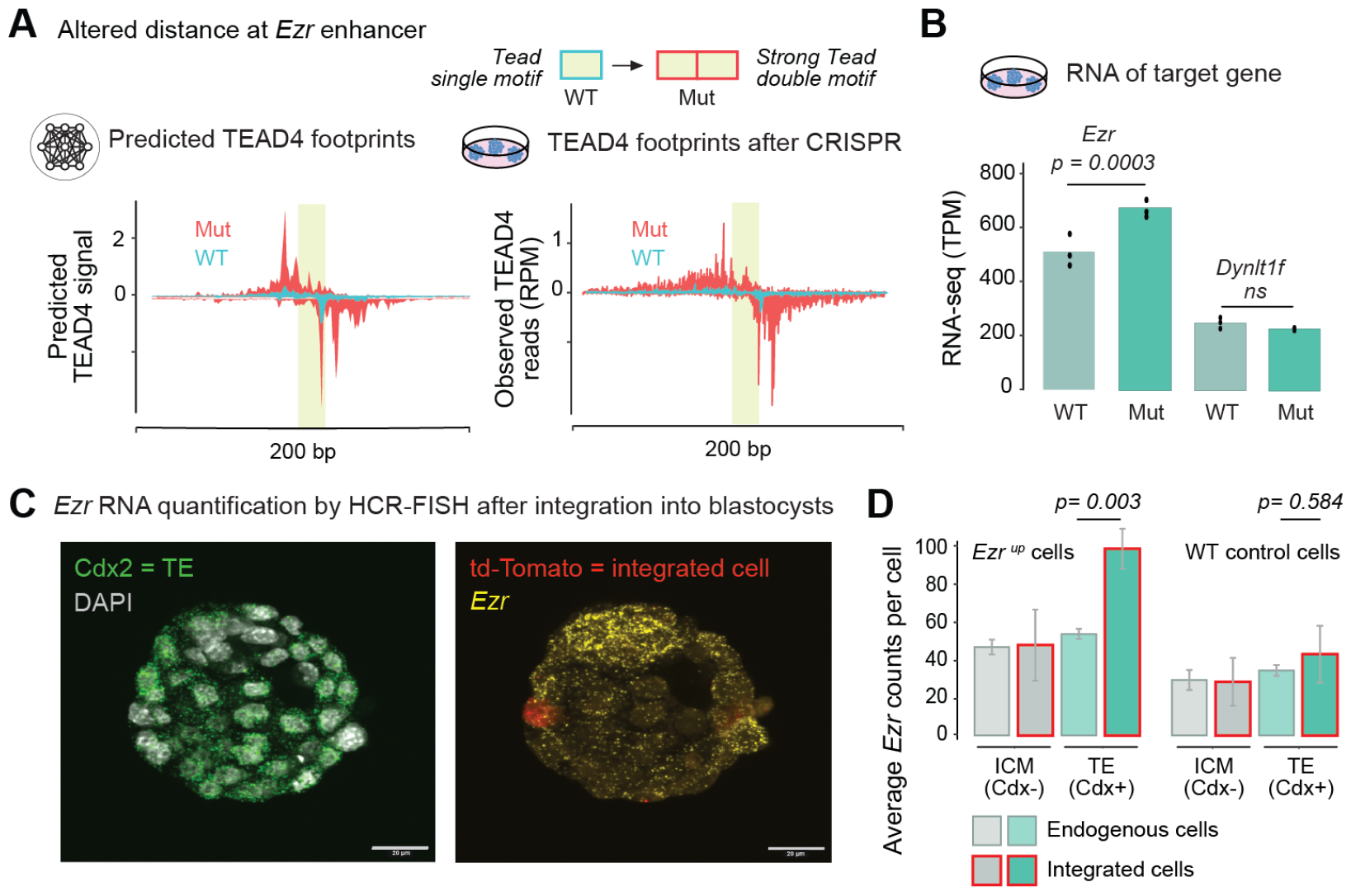
Enhancer design using BPNet and CRISPR-Cas9 increases target gene activity in mouse embryos. **A**) At the *Ezr* enhancer in mouse TSCs, the *Tead single* motif in the wild-type (WT) sequence was mutated (Mut) into a strong *Tead double* motif through CRISPR-Cas9 mediated homologous recombination. BPNet predicts increased TEAD4 binding (left), which was confirmed with ChIP-nexus experiments (right), normalized to reads per million (RPM). (mm10-chr17:6,827,705-6,827,905). The lime box represents the motif width. **B**) RNA-seq data normalized to transcript per million (TPM) in wild-type (WT) and mutant (Mut) cells. The p values for differential expression were derived using the edgeR package applied to three independent biological replicates. **C**) HCR-FISH was performed on aggregated mouse blastocyst embryos with wild-type (WT) or mutant (edited Ezr^up^) cells for probes *Cdx2, Ezr*, and *td-Tomato*. **D**) Quantification of average *Ezr* counts (average *Cdx2* counts shown in Figure S5E). *Ezr* transcripts were significantly increased for edited cells but not for wild-type cells among *Cdx2*+ trophectoderm (TE) lineage cells (student’s t-test *p* < 0.05).

We then used CRISPR-Cas9-induced homologous recombination to edit the endogenous enhancer in our TSCs (Figure S5A). We performed TEAD4 ChIP-nexus experiments on the edited cells and found that the TEAD4 binding footprint indeed matched the one predicted by BPNet (Figure 6A, right, and Figure S5B). We also tested the modified enhancer sequence in the luciferase assay and found it to increase enhancer activity compared to the wild-type sequence (Figure S5C). The increase was moderate (∼1.5-fold), likely because the wild-type enhancer activity was already high to begin with. To test whether this change affects the expression of *Ezr*, we performed RNA-seq on the edited cells with the *Tead double* motif and the wild-type cells with the *Tead single* motif. This revealed a moderate (1.5 fold), statistically significant increase in *Ezr* transcript levels in the edited cells (Figure 6B).

We next tested whether this edit increases *Ezr* expression in mouse embryos. We marked the edited TSCs (and wild-type TSCs as control) with Td-Tomato, aggregated these cells with early mouse embryos at the 4-8 cell stage, and analyzed the embryos when they reached the blastocyst stage, where the outer trophectoderm (TE) cells are clearly distinguishable from the inner cell mass (ICM) by their expression of *Cdx2*. We performed HCR-FISH to precisely quantify the expression of *Ezr* and *Cdx2* in these embryos (Figures 6C, S5D-F).

*Ezr* transcripts were specifically increased in edited cells but not wild-type cells, and only when they became TE cells with nuclear YAP1 (Figure 6C, right). Not all added TSCs maintained their TE identity but occasionally lost *Cdx2* expression and acquired ICM identity (Figure S5E), consistent with cell fate plasticity at this stage^43^. Notably, when the edited cells lost TE identity, *Ezr* transcripts were no longer increased. These findings show that the increased activity of the *Tead double* motif is specific to the cell type with active YAP1. They also demonstrate that, with the help of BPNet’s powerful predictive framework, enhancers can be manipulated toward cell type-specific activities and functions *in vivo*.

## Discussion

Here, we demonstrate how interpretable deep learning of high-resolution binding data identifies the cell-type-specific TFs that govern the transcriptional response of a signaling pathway. Using the Hippo pathway in mouse trophoblast stem cells as a model system, we show that the downstream component, nuclear YAP1, directly binds to TEAD4 as expected but that the other TF partners can be identified through the sequence rules that predict YAP1/TEAD4 binding levels. Since the deep learning model learns the rules *de novo* inside a “black box,” contributing TF motifs can be discovered even when the corresponding TF is not profiled. For example, in addition to TFAP2C, we found that AP-1 was also predicted to strongly contribute to YAP1 binding, consistent with evidence from other cell types^69–72^. The input from these partner TFs is predictive for enhancer activation and target gene expression and can be validated experimentally *in vivo*. Taken together, these results demonstrate that the target gene specificity of signaling pathways is read out by *cis*-regulatory sequences and that deep learning models are well suited to learn the complex sequence rules by which this occurs.

The extracted genome-wide sequence rules also provide insights into the mechanisms by which TFs boost the activity of signaling pathways. The syntax rules by which each TF motif enhances YAP1/TEAD4 binding levels as a function of distance fall into two categories: soft and strict motif syntax. This suggests that signaling pathways use at least two types of TF binding cooperativity. Both require DNA sequence as a template and require either no direct interactions or only very labile interactions between the cooperating TFs, which has implications for learning the rules of the *cis*-regulatory code across cell types. Our study also illustrates the power and limitations of using deep learning to derive the complex *cis*-regulatory rules that drive enhancer activation.

The first type of cooperativity by which signaling pathways receive input from cell type-specific TFs is through soft motif syntax, which we have characterized and validated for YAP1 and TFAP2C. In this type of TF cooperativity, binding enhancement occurs when the motifs are spaced within ∼150 bp and are strongest the closer the motifs are. Such sequence rules point to nucleosome-mediated cooperativity^15,20,26,117–121^ and may reflect the likelihood by which two motifs are covered by the same nucleosome. Although the mechanism is not well understood, it does not require specific interactions between TFs, which explains how signaling TFs can receive input from a wide variety of TFs in different cell types.

While nucleosome-mediated cooperativity is a flexible mechanism, we nevertheless found that some TFs had a much stronger contribution to YAP1 binding than others, suggesting that specific properties make them better partner TFs. The ability of TFs to bind and remove nucleosomes is such a key property^15,20^, but not all TFs that open chromatin necessarily help other TFs bind, and TFs could also contribute to YAP1 binding by activating transcription or forming condensates^20,122^. TFAP2C likely enhances TEAD4 binding by pioneering chromatin^123,124^, but it also interacts with co-activators to mediate activation^125^ and could help YAP1 form condensates^75,81^. Regardless of the mechanism, the input will likely also depend on the TF’s changing concentration over time, thus providing a means by which the input of TFs changes gradually during developmental transitions.

Although we set out to characterize the cell type-specific input to Hippo signaling, we also discovered a previously overlooked canonical response element of the pathway: *Tead double* motifs. These motifs use strict motif syntax and are much more widespread than previously observed. They are highly variable in sequence and mediate strong cooperativity even with many mismatches to the consensus; thus are an example of weak motifs with strong syntax^126^. As shown with MD simulations, this strong cooperativity results from DNA-mediated cooperativity, where the correctly spaced double motif brings two TEAD4 proteins in proximity to facilitate labile protein-protein interactions. We propose that such labile interactions are well suited for this type of cooperativity since they are strong enough to stabilize the complex but weak enough to be highly dependent on matching DNA sequences and thus read out the *cis*-regulatory code.

We note that this type of cooperativity is difficult to discover on a larger scale. Such labile protein-protein interactions cannot easily be predicted based on protein structures, and the sequence degeneracy of these motifs makes it challenging to discover them in the genome. This explains why the *Tead double* motif has occasionally been discovered in the past at low numbers yet has, as of now, not been considered a canonical element of the Hippo pathway. This type of cooperativity must, therefore, be discovered experimentally, either *in vitro*^85^ or through interpretable deep learning of genomics data, as we have done here. The improved discovery of TF motifs and syntax rules is, therefore, a key strength of this approach.

Another key strength of deep learning models is their predictive accuracy, which we leveraged here to design our experiments. Since BPNet accurately predicts the experimental outcome for sequences it has never seen during training, we can perform minimal mutations and predict the outcome of many experimental designs before choosing ones to pursue. In this manner, we changed a *Tead single* motif into a *Tead double* motif in an endogenous enhancer and showed that this increases the target gene expression in a cell type-specific manner *in vivo*. In the future, such focused sequence manipulations could be used to study and evaluate the potential impact of naturally occurring sequence variants and could help create enhancers with specific desired properties while minimizing side effects^6^.

A limitation of our approach is that it depends on high-quality binding data in the cell type of interest. While lower resolution data (ChIP-seq) can also be modeled by BPNet ^15,127^(see analysis of TEAD ENCODE data), it remains uncertain whether all relevant sequence rules were learned and extracted from the model. Even with the best data, careful analysis may reveal some limitations of the model. For example, we noticed that YAP1 binding correlates better

## Data and code availability

The raw and processed data for ChIP-nexus, ChIP-seq, ATAC-seq, TT-seq, and RNA-seq experiments have been deposited in GEO under series accession number GSE252463 and will be available following review. A web-accessible directory of the genomic datasets used in the paper can be viewed on the UCSC Genome Browser: Link.

The ChIP-nexus protocol and the data processing description can be found at https://research.stowers.org/zeitlingerlab/protocols.html.

The trained BPNet model will be available at Zenodo and Kipoi following review. Original data, including MD simulation trajectories and microscopy images, can be accessed from the Stowers Original Data Repository at http://www.stowers.org/research/publications/libpb-2440 following review. All code used to process and analyze the data can be accessed at https://github.com/zeitlingerlab/Dalal_hippo_signaling_2024.

## Acknowledgments

We thank Žiga Avsec, Robb Krumlauf, Helen McNeill, Anshul Kundaje, and Zeitlinger lab members for their helpful comments and suggestions on the manuscript. We thank the following Stowers Institute core facilities for their support: Sequencing and Discovery Genomics (Anoja Perera, Michael Peterson, and Amanda Lawlor), Histology (Dai Tsuchiya, Yongfu Wang, and Seth Malloy), Transgenic and Reproductive with enhancer activity markers than TEAD4 binding, yet the model interrogation suggested that YAP1 binding is mostly boosted through increased binding of TEAD4. Therefore, additional sequence rules may boost YAP1 binding, e.g., through enhancer-enhancer interactions^75,81^, which we did not model. We also did not model how enhancers regulate specific target genes, which is an important problem but beyond the scope of this study. In the future, it will be interesting to model how our derived enhancer rules are integrated into the larger genome context.

Technologies team (Michael Durnin and Andrea Moran), Cells Tissues and Organoids Center (Yan Wang, Naresh Kumar Rajendran, Sonia Ghosh, Maria Katt, Olga Kenzior, Shilpa Waduwawara, and Chongbei Zhao), Lab Services (Stacey Walker), Cytometry (Kevin Ferro, Jose Javier, KyeongMin Bae, and Jeff Haug), and Computational Biology (Hua Li, Madelaine Gogol, and Hassan Huzaifa). The research reported in this publication was supported by the Stowers Institute for Medical Research and NIH grant no. R01HG010211 to J.Z.

Author contributions: K.D. and J.Z. conceived the project as part of K.D.’s thesis research to fulfill the University of Kansas Medical Center requirements. K.D. and J.Z. designed the genomics and other experiments that K.D. performed. Deep learning model training, computational analysis, and *in silico* experiments were performed by K.D. and M.W. Additional genomics data analyses were done by K.D. ATAC-seq experiments were performed by S.K. MD experiments were conceived and designed by C.M., K.D, and J.Z., performed by C.M., and analyzed by C.M., K.D. and J.Z. Embryo aggregation, imaging, and analysis were conceived by K.D and J.Z. Imaging and analysis was performed by M.C.M. The manuscript was prepared by K.D. and J.Z. with input from all authors.

## Methods

### Mouse stem cell culture

Mouse trophoblast stem cells (TSCs) were a gift from Vijay Pratap Singh and were maintained in a feeder-free culture as described^54^. Briefly, feeder conditioned medium (Feeder-CM) was prepared by culturing γ-irradiated MEFs (mouse embryonic fibroblasts) in TS medium (RPMI 1640 medium, FBS 20%, 50 μg/mL of Penicillin and streptomycin (100×), 1 mM of Sodium pyruvate (100 mM), 0.1 mM of β-Mercaptoethanol (20 mM), 2 mM GlutaMAX (200 mM) for 72 h and then filtered with a 0.45 μm filter. 70% Feeder-CM plus 30% TS medium (70cond) supplemented with growth factor FGF4 (R&D System) and heparin (Sigma) (70cond + 1.5x F4H Medium) was used to maintain TSCs in feeder-free conditions. Mouse embryonic stem cells (ESCs) were cultured and maintained as previously described^15^.

### ChIP-nexus, PAtCh-Cap, and ChIP–seq experiments

For each ChIP-nexus experiment, 10e^6^ TSCs were used. Cells were washed with PBS and cross-linked with 1% formaldehyde (Fisher Scientific) in PBS for 10 min at RT. The reaction was quenched with 125 mM glycine. Fixed cells were washed twice with cold PBS and resuspended in cold lysis buffer (15 mM HEPES pH 7.5, 140 mM NaCl, 1 mM EDTA, 0.5 mM EGTA, 1% Triton X-100, 0.5% *N*-lauroylsarcosine, 0.1% sodium deoxycholate and 0.1% SDS), incubated for 10 min on ice and sonicated with a Bioruptor Pico (Diagenode) for five cycles of 30s on and 30s off. The ChIP-nexus procedure and data processing were performed as previously described^37^, except that the ChIP-nexus adapter mix contained four fixed barcodes (ACTG, CTGA, GACT, and TGAC), and PCR library amplification was performed directly after circularization of the purified DNA fragments (without the addition of the oligo and BamHI digestion). PAtCh-Cap was performed as previously described^128^ with 10% of sheared chromatin from 10e^6^ TSCs. ChIP-seq experiments, including a whole cell extract (WCE) control, were performed as described^129^ with 10e^6^ TSCs per ChIP. For each ChIP, 5-10 μg of antibody was coupled to 50-100 μl of Protein A or Protein G Dynabeads (Invitrogen). The following antibodies were used: anti-TFAP2C (R & D systems, no. AF5059), anti-TEAD4 (Abcam, no. ab58310), anti-CDX2 (Bethyl Laboratories, no. A300-692A), anti-GATA3 (Cell Signaling, no. 5852T), anti-YAP1 (Cell Signaling, no. 14074S), anti-Pol II (Cell Signaling, no. D8L4Y), anti-H3K27ac (ChIP-seq) (Active motif, no. 39135). For all experiments, at least two biological replicates were prepared–that is, the experiments were performed on different days, starting with cells from a different passage number. Single-end sequencing was performed on an Illumina NextSeq 500 instrument (75 cycles). The full ChIP-nexus protocol can be found on the Zeitlinger lab website at https://research.stowers.org/zeitlingerlab/protocols.html.

### Luciferase assays

Selected genomic regions of range 175-200 bp were synthesized using GeneArt Strings DNA fragments along with restriction enzyme sites for with Kpn1 (NEB, R0142) and XhoI (NEB, R0146) to clone in pNL3.2 vector. pNL3.2 vectors (Promega) were digested and cloned using an Infusion master mix (Takara) upstream of the luciferase gene. Stellar competent cells (Takara) were used for transformation and downstream miniprep (Qiagen), following the manufacturer’s protocol. The cloned sequences were confirmed using the Sanger sequencing method. 2.5e^5^ TSCs were used to transfect a total of 500 ng DNA with lipofectamine2000 in a ratio of 1:2 (DNA to lipofectamine2000), following the manufacturer’s protocol. Cells were co-transfected with 1:100 ratio of control (pGL4.54[luc2/TK]) and reporter construct (pNL3.2[NlucP/minP]). Cells were transfected in suspension for 15-20 min and resuspended with media to grow in each well of the 24-well plate.

Luciferase assays were performed using a Dual-Glo luciferase assay system (Promega). After 24h, cells were harvested, and the NanoDLR assay protocol was followed per the manufacturer’s instructions to take luminescence measurements with SpectraMax iD3 Plate Reader. Reporter luminescence signals were normalized according to their corresponding control luminescence signals, resulting in relative luciferase activity. Replicate luciferase assay experiments were performed independently three times (supplemental table 1 (sheet 2)).

### TT-seq experiments

TT-seq experiments were performed in three biological replicates on TSCs across three biological replicates, as described in^130^ https://www.protocols.io/view/transient-transcriptome-sequencing-experimental-pr-3byl42y22vo5/v1. Libraries were prepared using TruSeq Stranded Total RNA Library Prep Kit with Ribo-Zero Gold Set to degrade rRNA. Approximately 10e^6^ cells were seeded in a 100 cm dish (∼80% confluency) and incubated with 500 μM of 4sU (Sigma) at 37°C, 5% CO_2_ for 15 min. The cells were then collected by adding 4.5 ml of TRIzol lysis reagent (Life Technologies Corp), incubated for 5 min on ice, flash-frozen, and stored at −80°C. In the biotinylation of the 4sU labeled RNA step, acid-phenol-chloroform (ThermoFisher) was used instead of chloroform.

### ATAC-seq experiments

For each ATAC-seq experiment, 1e^5^ or 2e^5^ TSCs were harvested, washed with PBS, and resuspended in ATAC Resuspension Buffer (RSB, 10 mM Tris-HCl pH 8.0, 10 mM NaCl, 3mM MgCl_2_) with 0.1% IGEPAL CA-630. Tn5 transposition was performed as previously described^131,132^. Briefly, the cells were incubated for 3 min on ice in ATAC RSB supplemented with 0.1% IGEPAL CA-630, 0.1% Tween-20, and 0.01% Digitonin (Promega, G9441). The reaction was quenched with ATAC RSB with 0.1% Tween-20 and centrifugation. Tagmentation took place at 37°C and 1000 rpm for 30 min in a 50 μl reaction volume containing 10 μl of 5x Tagmentation Buffer (50 mM Tris-HCl pH 7.5, 25 mM MgCl_2_, 50% DMF), 0.5 μl 10% Tween-20, 0.5 μl 1% Digitonin, 1-2 μM assembled transposome and water. Tn5 transposase was purified in-house, as previously described^133^. Tn5 was loaded with previously reported oligonucleotides Tn5ME-A/Tn5mC1.1-A1block and Tn5ME-B/Tn5mC1.1-A1block^134,135^ by mixing equal amounts of purified Tn5 protein and annealed oligonucleotides for 30 min at RT. After tagmentation, the DNA fragments were purified using the Monarch PCR & DNA Cleanup Kit (NEB). Libraries were constructed using Illumina Nextera Dual Indexing, and qPCR was used to prevent over-amplification as described^131^. At least three biological replicates were generated, and paired-end sequencing was performed on an Illumina NextSeq 500 instrument (2 × 75 bp).

### RNA-seq experiments

TSCs wild-type and CRISPR-Cas9 edited cells were grown separately in wells of a 6-well plate and harvested at 80% confluency (∼2e^6^ cells) using 500 μl of TRIzol reagent (Life Technologies Corp). The lysate was incubated for 5 min on ice, flash-frozen, and stored at −80°C. For RNA extraction, the lysate was quickly thawed at 65°C, cooled on ice for 5 min, and vortexed. Then, 100 μl of chloroform was added per 0.5 mL of TRIzol lysis reagent, shaken vigorously for 15 sec, and incubated for 3 min. The sample was centrifuged at 4°C and 7000 x g for 25 min, and the upper colorless aqueous phase was transferred to a new tube. 250 μl of isopropanol was then added, incubated for 10 min at 4°C, and centrifuged at 4°C and 12,000 x g for 10 min. The total RNA precipitate formed a white gel-like pellet at the bottom of the tube, which was washed with 75% ethanol, air-dried for 5-10 min, and resuspended in 20 μl of RNase-free water. DNase treatment was performed using the TURBO DNase kit per the manufacturer’s instructions: adding 1 μl of TURBO DNase and 2 μl of DNase buffer to the dissolved RNA and incubating at 37°C for 30 min. To inactivate TURBO DNase, the RNA samples were extracted with phenol/chloroform (Sigma). The sample was centrifuged at 4°C and 7000 x g for 25 min, and the upper colorless aqueous phase was transferred to a new tube. 250 μl of isopropanol was added, incubated for 10 min at 4°C, and centrifuged at 4°C and 12,000 x g for 10 min. The total RNA precipitate formed a white gel-like pellet at the bottom of the tube, which was washed with 75% ethanol, air-dried for 5-10 min, and resuspended in 20 μl of RNase-free water. The samples were incubated in a water bath or heat block set at 55-60°C for 10-15 min. The RNA concentration was determined using a NanoDrop spectrophotometer, and the RNA integrity was checked on a 2100 Bioanalyzer using an Agilent RNA 6000 Nano Kit. mRNA-stranded libraries were prepared using a TruSeq poly-A Stranded mRNA Library Prep Kit and sequenced on an Illumina NextSeq 2000 P2 platform with 2 × 100 bp single-end reads. Three biological replicates were performed for wild-type and CRISPR-Cas9 edited cells.

### CRISPR-Cas9 experiments

In the first CRISPR TSC line, the *Tead double* motif on chr17:6,827,802-6,827,811 (mm10) was mutated from ACATTCCAGA (wild type) to GCATTCCAGGAATTCCA (mutant). In a second CRISPR TSC line, the *Tead single* motif (CACATTCCTA) on chr12:102,262,024-102,262,033 (mm10) was first inserted at 60 bp downstream from the *TFAP2C* motif (GGGCCCCAGGGCC) and then in a second round of CRISPR-Cas9 editing *Tead single* motif was mutated from CACATTCCTA (wild-type) to CACCGTCCTA (mutant) at its original position. crRNA target sites were designed using the IDT target predictor tool by evaluating the predicted on-target efficiency score and off-target potential. Alt-R CRISPR-Cas9 crRNA was designed to contain ∼40 bases of homology from the targeted cut site (gRNA and ssODN sequences are shown in supplemental table 1 (sheet 1)). Equimolar amounts (stock of 100 μm) of Alt-R crRNA and tracrRNA-ATTO550 were mixed to form gRNA at a final concentration of 50 μM. The mixture was heated at 95°C for 5 min and cooled at RT. The single-stranded donor oligonucleotides (ssODN) were designed to contain ∼40 bases of homology from the targeted cut site (crRNA and ssODN sequences were designed using the IDT software tool). A ribonucleoprotein (RNP) complex was formed by combining 150 pmol of gRNA (crRNA+tracrRNA) and 125 pmol of Cas9 HiFi v3 protein (IDT) with hybridization for 20 min at RT. The RNP was combined with 100 pmol of ssODN donor and 100 pmol of electroporation enhancer v2 and delivered to 1.5e^5^ cells by Neon electroporation (1,400 V, 10 ms, 3 pulses; Neon Transfection System, MPK5000, Life Technologies). Immediately after electroporation, cells were cultured in 0.5 μM Alt-R HDR enhancer V2 of 0.69 mM. After 24h, cells were washed with PBS before FACS sorting on S6 FACSymphony. Single cells were directly sorted into 96-well plates. Cells were screened for the expected mutations through paired-end sequencing on an Illumina MiSeq instrument (250 cycles). On-target indel frequency and expected mutations were analyzed using CRIS.py^136^. Clones with the intended homozygous mutation and sequence alignments >90% were chosen for further experiments, except for the 2nd CRISPR line, where we found one Indel and SNP within 500 bp of the original *Tead single* motif position, but these changes were predicted to be neutral by BPNet.

### Mice strains and superovulation

C57BL/6J (B6) strain of mice were used from the Stowers Institute for Medical Research (SIMR) core production colony. Three to four-week-old females were superovulated using 5IU of PMSG (Genway Biotech), followed by 5IU HCG (Sigma Aldrich) 46-48 h later. Following HCG, females were paired with B6 males and checked for a copulatory plug the next morning, indicating successful mating. Fertilized embryos were collected from the plugged females at 1.5 dpc (2-cell stage) by flushing M2 (Millipore) through the infundibulum and out the uterine horn using a blunt needle. Embryos were then incubated overnight at 37°C under 5% CO_2_ in humidified air in 4-well culture dishes containing KSOM media (Millipore). Experiments were approved by the SIMR IACUC and were performed following the committees’ guiding principles.

### Lentivirus transduction of fluorescent td-Tomato in TSCs

Two days (48 h) before the transduction of wild-type or CRISPR-Cas9 edited (putative *Ezr* region edits) TSCs, cells were seeded at 1 × 10^5^ cells per well in triplicate with 3 ml of media per well of 6-well plate. On the day of transduction, cells were small-sized colonies of about 30-40% confluence; the old media were removed and washed once with PBS and replaced with 2 ml of media. The cells were infected with prepackaged lentiviral particles (constitutive reporter vector expressing tdTomato fluorescent protein gene driven by EF1a promoter (Takara) at MOI of 20 (Stock: 3.5e^10^ TU/ml) along with polybrene (4ug/ml) (Sigma) for 24h before being replaced with a fresh medium. Four days after transduction, the td-Tomato-positive cells were selected using puromycin antibiotic selection (1μg/ml) (InvivoGen) and were kept under selection until the positive colonies reached 60-80% confluence. Once cells reached 80% confluence, the positive cells were FACS sorted on S6 FACSymphony, expanded, and used for embryo aggregation experiments.

### Aggregation assays to obtain chimeric embryos

To prepare the aggregation plates, six indentations on the bottom of the 35 × 10 mm plates were made using an aggregation needle (BLS) sterilized with 70% alcohol and added a drop of KSOM. All drops of KSOM were covered with mineral oil (Sigma). On the morning of the aggregation, the embryos (8-16 cell stage) were washed through M2 and then placed in drops of Tyrode’s solution (Sigma). After about 30 sec, the zona pellucida began to dissolve. Once the zona was dissolved, the embryos were picked up and rinsed through a drop of M2 to neutralize the Tyrode’s solution, then placed in a drop of KSOM. Embryos were moved from this dish to the aggregate plates, placing an embryo into each indentation. Clumps of td-Tomato transduced TSCs were then picked up with a mouth pipette and moved onto each embryo in the aggregate plate. Once settled and in contact with the embryo, the aggregation plates were cultured in an incubator at 37°C under 5% CO_2_ in humidified air for 46-48 h until the embryos reached the blastocyst stage. Chimeric blastocysts were fixed with 4% paraformaldehyde (ThermoFisher) for 20 min and washed three times with PBS before mounting them on a glass bottom plate (Cellvis) coated with poly-L-lysine (Sigma). Embryos were imaged with a Zeiss LSM800, an upright confocal laser scanning microscope.

### Immunofluorescence stainings of chimeric embryos

A few fixed chimeric blastocysts were used for immunofluorescence stainings. The embryos were washed three times with PBS-T (PBS with 0.1% Triton X-100) and incubated in PBS-T for one hour or longer at 4°C. Embryos were then washed two times with PBS for 10 min each and incubated with 300μl of superblock solution (ThermoFisher) for 90 min at RT, before adding the primary antibodies: CDX2 (BioGenex-MU392A-5UC) and Nanog (Cell Signaling, 8822S) with 10 μg/ml of DAPI from BioLegend:422801. The CDX2 antibody came with a signal enhancing reagent, which was used to replace 75% of the superblock solution while incubating with the primary antibody. The embryos were incubated overnight at 4°C, covered from light. The next day, the samples were washed for 10 min each three times with PBST (0.1% Triton X-100) at RT and once with PBS at RT. Secondary antibodies were added (biotium:20015,20047) in special PBS (ThermoFisher) at a 1:300 dilution with DAPI 2 μl in 1 ml of 10 μg/ml (BioLegend) and kept on light rotation for 2 h at RT. Samples were then washed three times with PBS for 10 min. Samples were imaged immediately or kept for up to a week at 4°C before imaging. Imaging was performed with an upright confocal laser scanning microscope (Zeiss LSM800) with 40x magnification. Maximum intensity Z projections and adjustments to the brightness and contrast were performed in ImageJ/FIJI ^137^. Samples larger than the field of view were taken as tiled images and stitched with the Grid/Collection Stitching plugin in ImageJ.

### HCR-FISH on chimeric embryos

Embryos were fixed in 4% paraformaldehyde for 20 min and washed three times in PBS + 0.2% Triton-X for 10 min each. RNA FISH experiments were performed using HCR v3.0 (Molecular Instrument Inc.). The RNA sequences that were used to design probes are listed along with the chosen amplifiers and probe set size: Ezr (NM_009510.2, B4,32), CDX2 (NM_007673.3, B1,29), tdTomato (B5,16). The following amplifiers with Alexa fluorophore were used: 488, 546, and 647. The fixed embryos were serially dehydrated into methanol and stored at −20°C until use. To rehydrate the embryos, they were washed in PBS + 0.1% Triton-X (PBST). Embryos were incubated in the hybridization buffer for 30 min at 37°C, then in the hybridization buffer containing the probes at 37°C for 16h. Embryos were washed 5 times with the wash buffer for 5 min each, then 2 times in 5x SSCT (5x SSC + 0.1% Tween 20). Amplifiers were snap-cooled by heating at 95°C for 90s and cooled to RT for 30 min under dark conditions. Embryos were incubated in an amplification buffer for 30 min at RT before adding the amplifiers and incubating the embryos for 80 min at RT in a humid chamber under dark conditions. Embryos were washed 4 times in 5x SSCT for 5 min each, stained with DAPI (10 ug/ml) from BioLegend in 5x SSCT for 30 min, and then washed two times in 5x SSCT. Embryos were stored in 5x SSC at 4°C until imaging. Images of labeled chimeric blastocysts were acquired with an Orca Flash 4.0 sCMOS at full resolution on a Nikon Eclipse Ti2 microscope equipped with a Yokagawa CSU W1 Spinning Disk Confocal with 50 μm pinholes. A Nikon 40x long working distance water immersion objective, NA 1.15, was used to acquire all channels with exposure times of DAPI: 20ms, Alexa488: 200ms, Alexa546: 250ms, and Alexa647: 250ms.

### Molecular dynamics simulations

System preparation and simulation procedure: Canonical B-form DNA was created for each simulated sequence using Avogadro 1.2.0^138^. The TEAD4 structure was taken from PDB 5GZB^111^, and selenomethionine residues were replaced with regular methionine by simply renaming the selenium atom to sulfur. In order to align the protein structure from 5GZB onto the created DNA structures, we aligned the phosphorus atoms from the 4th to 10th residue on chain B of the PDB (which correspond to the bases CATTCCT) to the corresponding atoms on the created DNA. Since we simulated TEAD4 dimers, we performed this alignment twice, once for each binding site. This alignment was accomplished using VMD 1.9.3.^139^. We combined the two translated copies of the TEAD4 protein and the synthetic DNA sequence into one system using AmberTools20 (Case 2020). We used the ff19SB force field for protein atoms^140^, the bsc1 force field for DNA^141^, and the OPC for water and ions^142^. Systems were solvated in truncated octahedra of water with a 12 Å padding between the solute and cell edge. Systems were charge-neutralized with K+ ions, and additional K+ and Cl-ions were added to bring the system to a concentration of approximately 150 mM KCl. Systems were minimized using NAMD 2.13^143^. During minimization, a cutoff distance of 9 Å was used, and solvent bonds were held rigid, though all solute bonds were unrestrained. A timestep of 1 fs was used, and PME electrostatics was applied with a grid spacing of 1 Å. Ten thousand steps of minimization were performed. For thermalization, we used a GPU-enabled build of NAMD 2.14^143^. The same parameters were used as in the minimization, except for the introduction of a Langevin piston to maintain the system pressure at 1 atm and a harmonic collective variable restraint^144^ to prevent the ends of the DNA from fraying during the simulation. This restraint was applied between H1 from the terminal guanine and N3 of the terminal cytosine. (The DNA ends with a GC pair on each end, and both ends of the DNA were restrained in the same way.) A force constant of 1 kcal/mol/Å^2^ was applied to maintain this distance at 2 Å. During thermalization, all velocities were started from zero and gradually warmed by applying a Langevin thermostat to raise the system temperature to 310 K during a ten ps simulation. The thermalized systems were equilibrated for ten ns, the only difference in configuration from the thermalization simulation being the timestep (increased from 1 fs to 2 fs) and the use of rigid bonds (all bonds, including hydrogen, was made rigid during equilibration and production. Coordinates were saved for every ps for both equilibration and production runs. We have provided dehydrated trajectories along with all analysis scripts in Python (Python.org), D (dlang.org), and VMD^139^ in one folder, which will be provided upon peer-reviewed publication of this work or on request. Complete, hydrated trajectories, totaling approximately 5 TB of data, are available upon reasonable request.

### ChIP-nexus data processing

ChIP-nexus and PAtCh–Cap single-end sequencing reads were pre-processed by trimming off fixed and random barcodes and reassigning them to FASTQ read names. ChIP-nexus adapter fragments were trimmed from the 3’ end of the fragments using cutadapt (v.2.5)^145^. ChIP-nexus and PAtCh–Cap reads were aligned using bowtie (v.1.1.2)^146^ and its bowtie to the *Mus Musculus* genome assembly mm10. Aligned ChIP-nexus and PAtCh–Cap BAM files were deduplicated based on unique fragment coordinates and barcode assignments. ChIP-nexus coverage was normalized was acquired through reads per million (RPM) normalization, where the ChIP-nexus sample coverage was scaled by the total number of reads divided by 10^6^. ChIP-nexus peaks were mapped using MACS2(v.2.2.6)^147^ with parameters designed to restimulate the full fragment length coverage instead of the single stop base coverage (--keep-dup=all-f=BAM--shift=-75--extsize=150). ChIP-nexus peaks were filtered for reproducibility in a pairwise fashion using the Irreproducible Discovery Rate framework (IDR) (v.2.0.3)^148^. The IDR framework selected the peaks used for downstream analysis from the largest pairwise comparison.

### ChIP-seq data processing

ChIP-seq single-end sequencing reads were aligned to the *Mus Musculus* genome assembly mm10 using bowtie2 (v.2.4.2)^146^. Aligned ChIP-seq BAM files were deduplicated based on unique fragment coordinates and fragments extended based on the average experiment fragment length as determined with an Agilent 2100 Bioanalyzer. Normalized ChIP-seq coverage was acquired using the deepTools subfeature bamCompare (v.3.1.3)^149^ using parameters to generate RPKM or log_2_ fold-change scaling (--scaleFactorsMethod=None –normalizeUsingRPKM--binSize=50) or (--scaleFactorsMethod=readCount--operation=log2--binSize=50). ChIP-seq peaks were mapped using MACS2 (v.2.2.6)^147^ with default parameters and an applied background coverage using the associated WCE ChIP-seq control experiment. ChIP-seq peaks were filtered for pairwise reproducibility using the Irreproducible Discovery Rate framework (IDR) (v.2.0.3)^148^.

### TT-seq data processing

TT-seq 75 bp paired-end sequencing reads were aligned using STAR(v.2.7.3)^150^ to the *Mus Musculus* genome assembly mm10 with the following parameters: outFilterMismatchNmax 2, outFilterMultimapScoreRange 0. SAMtools (v.1.14)^151^ were then used to keep alignments with mapping quality greater than 255 (-q 255), and only proper pairs (-f 2) were selected. Strand-specific BAM files for each replicate and combined were generated using the following parameters (samtools view -b -f 128 -F 16; -b -f 80; -b -f 144; -b -f 64 -F 16) and (samtools merge plus_128.bam with plus_80.bam and minus_144.bam with minus_64.bam). Normalized TT-seq coverage was generated using bamCoverage (v.3.1.3)^152^ parameter Reads Per Kilobase per Million mapped reads (RPKM).

### ATAC-seq data processing

ATAC-seq paired-end sequencing reads were aligned using bowtie2 (v.2.3.5.1)^146^ to the *Mus Musculus* genome assembly mm10. Normalized ATAC-seq coverage was acquired through RPKM normalization along with following parameters: -bs=50 --minFragmentLength 10 --maxFragmentLength 1000 --ignoreDuplicate --extendReads

### RNA-seq data processing

RNA-seq 100 bp single-end sequencing reads were aligned to the *Mus Musculus* genome assembly mm10 using STAR (v.2.7.3)^150^ with the following parameters: outSAMtype BAM SortedByCoordinate, outSAMprimaryFlag OneBestScore, outFilterMultimapNmax 20, outFilterMismatchNoverLmax 0.1, outFilterType BySJout, alignSJoverhangMin 8, alignSJDBoverhangMin 1,outFilterMismatchNmax 999,alignIntronMin 20,alignIntronMax 1000000,alignMatesGapMax 1000000,limitBAMsortRAM 10000000000, outSAMattributes NH HI MD AS nM, quantMode TranscriptomeSAM GeneCounts. Rsem-calculate-expression (v1.3.0)^153^ was used to generate an expression table with the following parameters: no-bam-output, estimate-rspd, strandedness reverse.

### HCR-FISH image analysis

Images were analyzed in Python 3.9. 3D masks were created with the DAPI label using the Cellpose deep learning package^154^. After a small Gaussian blur of width 1×2×2 pixels, Cellpose segmentation was performed with the cyto model with a diameter of 60 and minimum cell size of 10000. The trophoblast cells were segmented well with this method in 3D, but the crowded inner cell mass cells were frequently corrected by hand using Napari (doi:10.5281/zenodo.3555620). The masked DAPI was expanded by 4 pixels in the xy direction to encompass more of the cytoplasm HCR label in each cell. The small HCR puncta were found by first performing a gaussian blur of 2×5×5 width, then a Laplace filter using Gaussian derivatives with sigma = 0.1, 0.5, 0.5. Finally, the local maximum peaks in intensity are found using the Scikit-image peak_local_max function with a threshold of 11 for the CDX2 channel and 15 for both Ezrin and tdTomato channels. The number of HCR puncta found inside each masked cell was recorded. A threshold was determined to categorize a cell as CDX2 positive (greater than 5 HCR puncta) or tdTomato positive (greater than 15 HCR puncta). The threshold for tdTomato is greater because the HCR hairpin signal is sometimes found on the outside surface of the blastocyst, forming brighter and larger puncta compared to the interior cell signal, which would cause too many cells to be categorized as tdTomato positive. All thresholds are held constant between all images of blastocysts.

### Molecular dynamics analysis and visualization

Solvent-accessible surface areas were calculated using VMD^139^, with a 1.5 Å radius around all atoms. The buried surface area between the two systems was calculated by subtracting the surface area of the combined system from the sum of the surface areas of each component system. Plots were generated using Matplotlib^155^, Scipy^156^, and NumPy^157^ with Python 3.10 (python.org). Figures were generated with Tachyon^143^ in VMD; trajectory frames were aligned using a frame-by-frame aligner developed previously^158^. Secondary structures were determined using STRIDE^159^.

### BPNet model training

BPNet (v.0.0.23) architecture and software were applied as previously described^15^. Model inputs were 1000 bp genomic sequences centered on the ChIP-nexus peaks of TF of interest. Model outputs were the predicted counts (total reads across each region) and predicted profile (coverage signal across each region) for TFAP2C, TEAD4, CDX2, YAP1, and GATA3 ChIP-nexus experiments. ∼150K IDR-reproducible peaks from TFAP2C, TEAD4, CDX2, YAP1, and GATA3 ChIP-nexus experiments were pooled and used as model inputs. Validation datasets were peaks across chr5,6,7,19; test datasets were peaks across chr1,8,9, and peaks across chrX and Y chromosomes were excluded from the analysis. The remaining regions were used for model training. Hyper-parameters were the default BPNet architecture. The trained model performance was assessed by comparing (1) area under the PrecisionRecall Curves (auPRC) for profiles over different bins of resolution between observed ChIP-nexus profiles and predicted BPNet profiles (Figure S1C) and (2) counts correlations of observed ChIP-nexus signals to predicted BPNet signals for each TF (Figure S1D) as previously described^15^. The auPRC values were benchmarked alongside replicate-replicate, observed random, and observed-average observed profile comparisons to establish an in-context understanding of predicted profile accuracy. All BPNet models were implemented and trained using Keras (v2.2.4), TensorFlow1 backend (v.1.70), and the Adam optimizer150. The training used an NVIDIA® TITAN RTX GPU with CUDA v9.0 and cuDNN v7.0.5 drivers. To obtain the *Tead double* motifs in ESCs for analysis Figure 5G, TEAD4 ChIP-nexus experiments were pooled and used as model inputs to train a single TF model; ∼15K IDR-reproducible peaks were used. Validation peak datasets across chr 1,7,3,14, test peak datasets across chr2,8,9, and peaks across chromosomes X and Y were excluded from the analysis. Hyper-parameters, model performances, and BPNet implementation were performed as described above. PAtCh-Cap control in ESCs was from^15^. We performed DeepLIFT and TF-MoDISco on the trained model to generate an ESCs-specific *Tead motif* set. For analysis in Figure 5G, we used *Tead double* motifs from fold 1. Additional models were trained with the same architecture as part of three-fold validation (fold 2 and fold 3). Spearman counts correlation values (top right) were determined by comparing the observed ChIP-nexus counts with BPNet’s predicted counts at TEAD4 ChIP-nexus peaks in ESCs (Figure S5L).

### Motif extraction, curation, and island generation

DeepLIFT (v0.6.9.0, derived from the Kundaje Lab fork of DeepExplain (https://github.com/kundajelab/DeepExplain)^58^ was applied to the trained BPNet model to generate the contribution of each base across a given input sequence to the predicted output counts and profile signals. Contribution scores for counts and profile outputs were generated for all 5 TF tasks. TF-MoDISco (v0.4.2.2)^59^ was then applied for each TF separately. Regions of high counts contribution were identified, clustered based on within-group contribution and sequence similarity, and then consolidated into motifs. The *Tfap2c, Tead, Cdx2, Gata3, and Yap1* motifs were manually identified based on their similarity to the known motif and the sharp average ChIP-nexus binding footprint of the corresponding TF. Once motifs were characterized and confirmed, they were used to label genomic instances by CWM scanning as previously described^15^. Briefly, a motif was mapped based on both Jaccardian similarity to the TF-MoDISco contribution weight matrix (CWM) and sufficient total absolute contribution across the mapped motif. Then, motifs were filtered for redundant assignment of palindromic sequences and overlapping peaks. To obtain regions of mapped motif combinations with enhancers for downstream measurement of enhancer activity to get specific mapped motif pairs, ‘motif islands’ were generated as described^20^. Each island starts as a 500 bp (enhancer window) region centered on the motif and gets clustered and merged with another nearby motif island if they overlap. In this manner, islands get extended if a motif is within less than 500 bp. The motif islands, by their motif combinations with motif numbers, read sums of TFs binding and enhancer activity (provided in supplemental table 1 (sheet 3)).

### Visualization of YAP1 binding and enhancer activity markers

To visualize the correlation between YAP1 binding and the markers of enhancer activity, we selected regions using the following criteria: BPNet-mapped motifs that were absent of ERVs, were within TEAD4 peaks, and showed TEAD4 binding. At those regions, we calculated the total ChIP-nexus read counts for YAP1, selected regions above the median value, and sorted based on the total read counts. These regions were then divided into the top 5000 regions with high YAP1 reads and the 5000 regions with median YAP1 binding. We used this set to calculate normalized reads and generate the TEAD4, YAP1, H3K27ac, Pol II, and Nascent-RNA heatmap.

### Motif pair interaction analysis

We selected mapped regions with only one motif pair from the motif islands set for the following motif-pair combinations: *Tfap2c-Tead, Cdx2-Tead, Gata3-Tead4, and Tfap2c-Tead double*. We then sorted the regions by the distance between the two motifs and included distances less than 160 bp for display. YAP1 contribution scores from the binding model were used to make heatmaps in ggplot. The *in silico* motif interaction analysis and odds ratio calculations for the co-occurrence likelihood of motif pairs were performed as described^15^.

### Enhancer regions selection for reporter assay

This analysis was to predict TFAP2C and TEAD4 binding on genomic regions with different motif distances and how this changes upon editing the distance between the motifs. From our islands, we selected regions with one Tead single and one *Tfap2c* motif within less than 200 bp and resized them to 400 bp putative enhancers, and recorded the coordinates of *Tfap2c* and *Tead4* motifs within the putative enhancers for mutations. We then identified the nearest genes using the biomaRt package. For each putative enhancer, we generated sequences for wild-type, mutated motif for each at its original position by mutating the two most contributing nucleotides to the least contributing within that motif. Then, we inserted the same motif at distances in multiples of 10 or 15 within a 400 bp window. These sequences were combined into an array to predict TF binding and contributions at a motif range of 50 bp. The resulting unique enhancer values were exported in R for plotting. The luciferase assay and CRISPR regions were selected by high binding of TEAD4, TFAP2C, and YAP1 at these putative enhancers and by context-relevant gene targets.

### Extracting regions with different *Tead double* motif spacings

To map regions in the mm10 genome with different *Tead double* motif spacings, we used pattern matching (with no mismatches) to identify instances of a single Tead motif (RMATTCCWD). Then, regions with two motifs within 23 bp were identified, and the frequencies by which each motif spacing occurred were recorded. Thus, for a motif spacing of 2, the matched sequence is RMATTCCNNRMATTCCNN. The predicted TEAD4 binding signal was then calculated for all motifs injected into randomized sequences and averaged over 256 iterations. The results from each spacing were then averaged.

### *Tead* motifs variant analysis

To assess the distribution of motif variant frequency, identical sequence patterns of CWM-mapped *Tead single* and *double* motif patterns were grouped, analyzed and visualized. To obtain a robust representation, only patterns that occurred in the top 90th percentile and occurred at least 10 times were considered. After injecting each sequence pattern into 256 random sequences, BPNet was used to predict TEAD4 binding. The average predicted signal for each pattern, along with the pattern frequency, was plotted using ggplot.

### Genome-wide TEAD4 binding cooperativity on *Tead double* motifs

This analysis aimed to investigate the potential synergy between each side (each *Tead* motif) of the *Tead* double motifs. We selected regions that did not overlap with either ERVs or promoter regions, extracted the sequences of the *Tead double* motifs, and oriented them in the 5’>3’ direction. We then split the motif sequences into two half-sites, each corresponding to a *Tead single* motif. We then predicted the binding of TEAD4 at the half-sites and the complete double motifs injected into random sequences. The values for the two half-sites were summed and compared to those for the complete double motifs as a measure of synergy between the two half-sites of TEAD4 double motifs.

## Supplementary Figures

**Figure S1.**
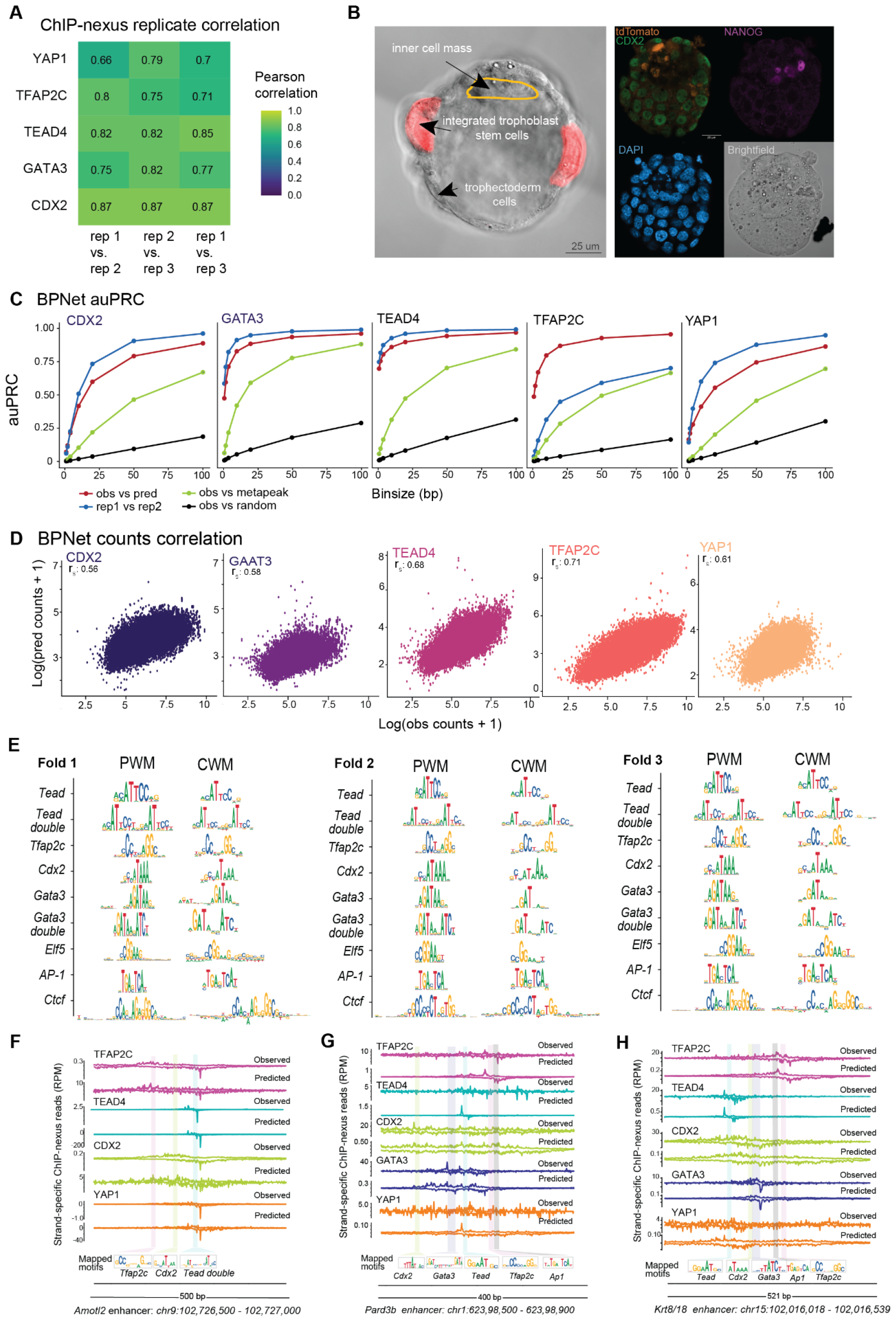
BPNet accurately learns the profile and counts information for TFs important in TSCs (related to Figure 1) **A**) All pairwise comparisons Pearson correlation values of TF ChIP-nexus experiments between replicates. The coverage for each replicate was calculated across a 200 bp window centered on the MACS2-called peaks for each TF. Because ChIP-nexus provides strand-specific information, the absolute value of the counts from the negative strand, which would otherwise be negative, was taken and added to the counts across the positive strand to determine the total region counts for a given replicate. **B**) Upon performing aggregation assay at the blastocyst stage of mouse embryos, integrated TSCs expressing td-Tomato lentivirus construct get integrated with the outer trophectoderm (TE) layer cells, thus closely resembling the fate of the neighboring cells from where they were derived (left). Immunofluorescence staining on an aggregated embryo reveals an overlap between CDX2, a marker for the trophectoderm layer, and td-Tomato cells, in contrast to NANOG, a marker for the inner cell mass, suggests a preference for TSCs to aggregate with TE cells (right). **C**) Area under the Precision-Recall Curves (auPRC) shows that BPNet accurately predicts the profile positions. The ability of BPNet to identify positions of high ChIP-nexus signal is assessed at various resolutions up to 100 bp. Replicate experiments, average ChIP-nexus profiles, and randomized profiles are shown as controls. **D**) BPNet predicts ChIP-nexus counts with high accuracy. Spearman counts correlation values were determined by comparing the observed ChIP-nexus counts with BPNet’s predicted counts at ChIP-nexus peaks for each TF of interest. **E**) The representative short motifs discovered with TF-MoDISco contain known motifs, motifs for TF we had not profiled, and known motifs new in this context. Additional models trained with the same architecture returned the same set of motifs as part of three-fold validation (fold 2 and fold 3). All sequence logos share the same *y-axis*. **F-H**) Comparing experimentally generated TF binding with BPNet-predicted TF binding at the putative *Amotl2* enhancer (F), *Pard3b* enhancer (G), and *Krt8/18* enhancer (H) illustrates BPNet’s predictive accuracy. Each color is a different TF, where the top track is the experimental ChIP-nexus data, and the bottom track is the predicted binding. Motifs were identified and mapped by BPNet. Putative *Amotl2* and *Pard3b* enhancers were on the withheld chromosome during BPNet training. The putative *Krt8/18* enhancer shows predictive accuracy of the fuzzy profile of YAP1, which is an indirect binder.

**Figure S2.**
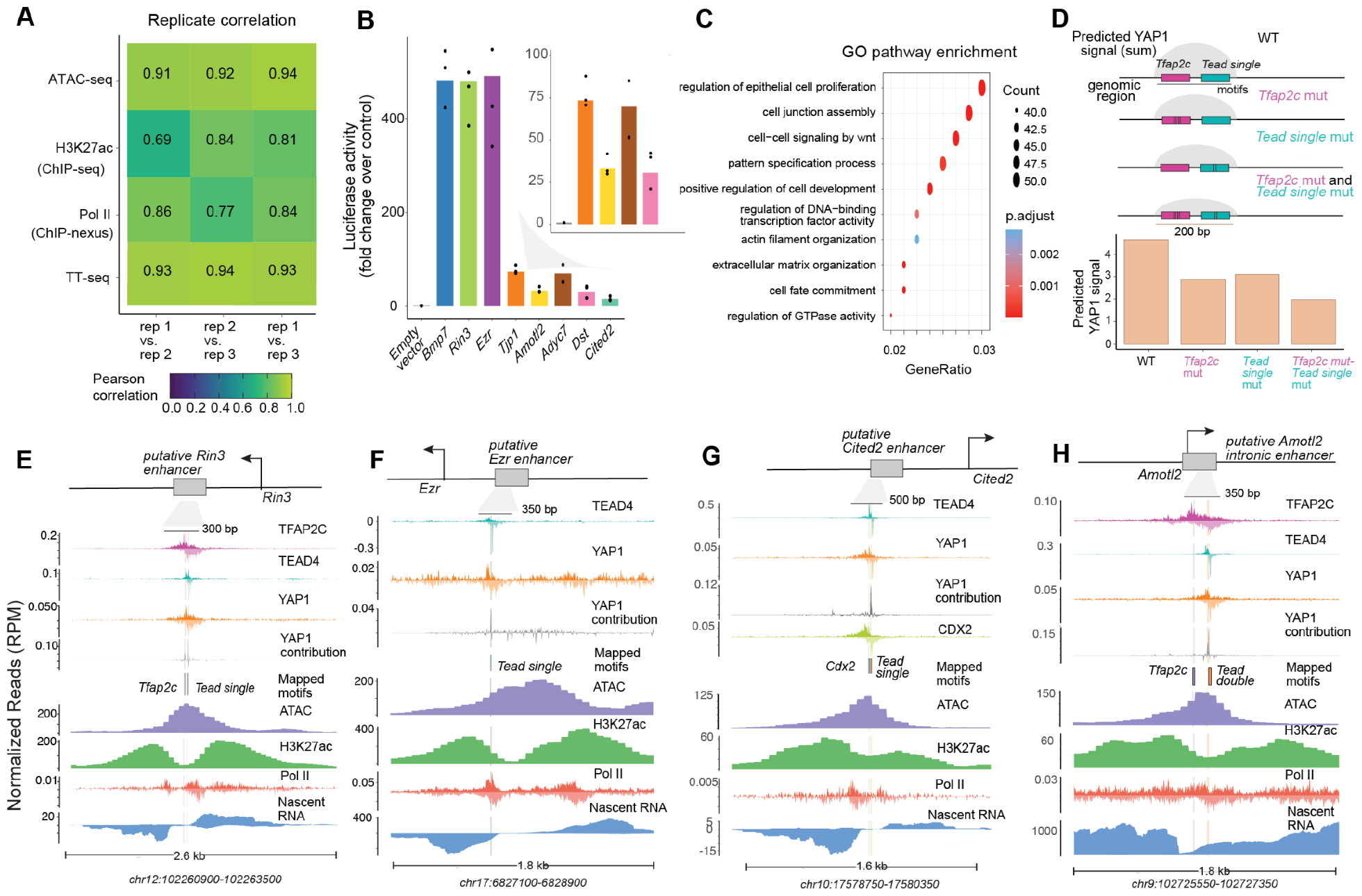
TSCs specific enhancers show activity markers with TFs-bound motifs. (related to Figures 2 and 3) **A**) Pearson correlation values were determined for all pairwise comparisons between replicates of ATAC-seq, H3K27ac ChIP-seq, Pol II ChIP-nexus, and TT-seq experiments. For Pol II ChIP-nexus, coverage for each replicate was calculated across a 200 bp window centered on the MACS2-called peaks. Because ChIP-nexus provides strand-specific information, the absolute value of the counts from the negative strand, which would otherwise be negative, was taken and added to the counts across the positive strand to determine the total region counts for a given replicate. For ATAC-seq, counts for each replicate were calculated across a 600 bp window centered on the MACS2-called peaks. For ChIP-seq, counts for each replicate were calculated across a 1000 bp window centered on the MACS2-called peaks. For TT-seq, counts for each replicate were calculated across a 500 bp window centered on the MACS2-called peaks of Pol II. **B**) Luciferase assay of the wild-type 175 bp or 200 bp minimal putative enhancers consisting of either the *Tead single* and *Tfap2c* motif pair or the *Tead double motif* was performed thrice as biological replicates and normalized over the empty vector control. **C**) The *Tead single* and *Tfap2c* motif-pair islands mapped within 160 bp distance were used to find the nearest gene and perform gene ontology analysis using the clusterProfiler package. **D**) Average YAP1 predicted signal summed across 200 bp window (as portrayed in grey color in the graphic) of putative *Bmp7* enhancer with mapped *Tead single* and *Tfap2c* motif in wild-type, individual motif mutated, and both motif mutated sequences. **E-H**) The genome track of putative active enhancers for genes E) *Rin3*, F) *Ezr*, G) *Cited2*, H) *Amotl2* with mapped motifs, normalized ChIP-nexus TFs binding profiles, and normalized read pileups of enhancer activity markers.

**Figure S3.**
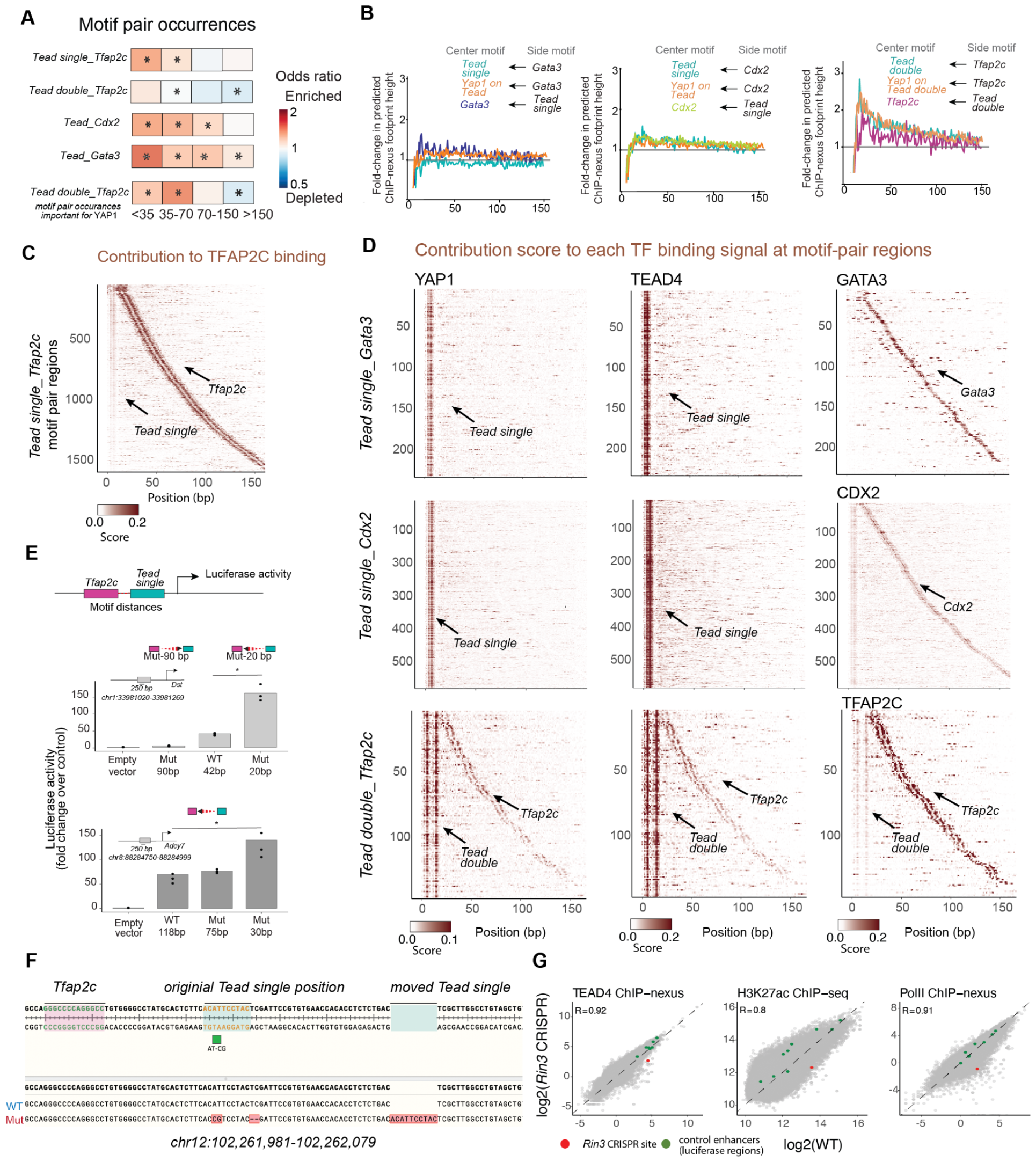
The distance-dependent cooperativity of TEAD4 with TFAP2C is motif-specific and directional (related to Figure 3). **A**) Mapped motifs enriched at short distances, calculated as the odds ratio of the frequencies observed for wild-type over permuted regions. Significance was denoted by \**p* < 10^−5^ using Pearson’s chi-squared test. **B**) In the *in silico* analysis, motifs are injected into randomized sequences, and BPNet is used to predict the average enhancement of TF binding to its motif (center) in the presence of a side motif^15^. The results show no distance-dependent TEAD4 and YAP1 binding enhancement in the presence of *Gata3* or *Cdx2* motifs. We see mutual binding enhancement of TEAD4, YAP1, and TFAP2C at a closed range distance with *the Tead double-Tfap2c* motif pair. **C**) Heatmap showing BPNet contribution scores of TFAP2C binding across regions with one *Tead single* and one *Tfap2c* motif, ordered by the distance between the motifs (up to 160 bp). There is no visually strong contribution from the *Tead single* motif. **D**) Heatmap showing BPNet contribution scores of TFs across motif-pair islands of *Tead single-Gata3, Tead single-Cdx2, and Tead double-Tfap2c*, ordered by the distance between the motifs (up to 160 bp). **E**) Luciferase assay of the wild type and mutated 200 bp minimal putative enhancer of *Dst (*mm10*-chr1:33981020-33981269)* and *Adcy7* (mm10-chr8:88284750-88284999) were performed thrice as biological replicates and normalized over the empty vector control. Significance was determined by a student’s t-test (*p* < 0.05). Increasing the distance between *Tead single* and *Tfap2c* decreases activity and vice-versa. **F**) Sanger sequencing chromatogram at the putative *Rin3* enhancer with mapped and moved motifs, where the wild-type sequence has a distance of 20 bp between *Tfap2c* and *Tead single* motif. The sequential CRISPR, where the *Tead single* motif was inserted away from the *Tfap2c* motif to generate mutant cells with a new distance of 60 bp between *Tfap2c* and *the Tead single* motif. Then, the most important nucleotide (highlighted in the green box) within the *Tead single* motif at the original position was mutated to abolish TEAD4 binding. **G**) Pairwise comparisons between WT and CRISPR clone cells show high Pearson correlations for TEAD4 and Pol II ChIP-nexus and H3K27ac ChIP-seq for control enhancers (green dots) from the luciferase assay (Figure S2B) and the selected *Rin3* enhancer (CRISPR site in red dot) were consistent between the two replicates.

**Figure S4.**
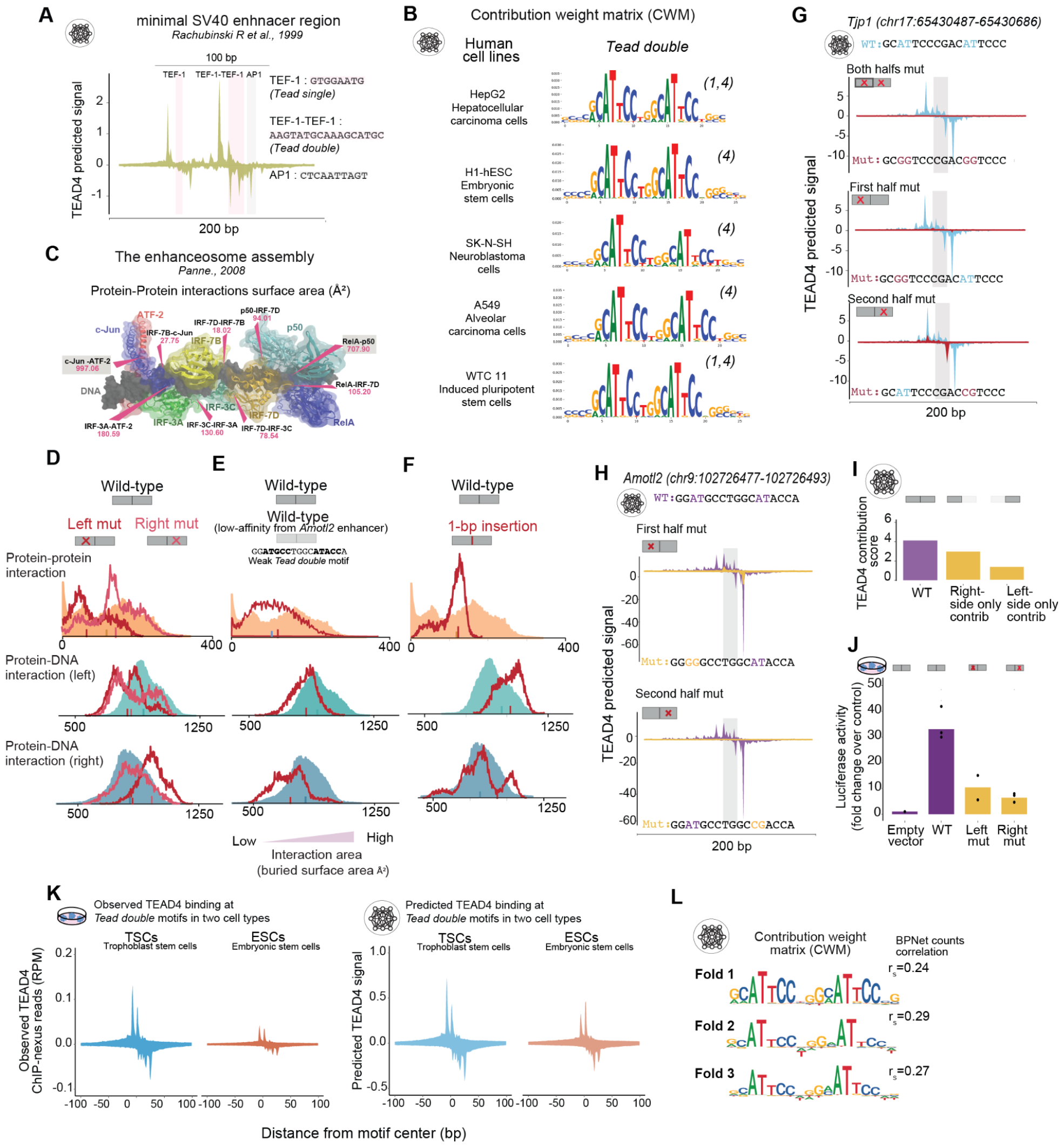
BPNet and MD simulations reveal quantitative details of TEAD4 cooperative binding at *Tead double* motifs (related to Figures 4 and 5) **A**) BPNet predicted TEAD4 ChIP-nexus footprint at minimal SV40 enhancer (100 bp) consisting of TEF-1 (*Tead single*) and TEF-1-TEF-1 (*Tead double*) motif sequence patterns. **B**) The contribution weight matrix (CWM) of *Tead double* motifs of either human TEAD4, TEAD1, or both (shown in brackets) when trained as a single TF model in various human cell lines obtained from ENCODE Consortium highlights the nucleotide contribution to TEAD binding predictions. We show here data generated by the lab of Richard Myers, HAIB, with the following identifiers: ENCSR934WOF; ENCSR497JLX, ENCSR285HHZ, ENCSR800JRG, ENCSR000BUQ, ENCSR000BRY, ENCSR000BUD and deep learning model trained by Anshul Kundaje’s lab. **C**) The predicted enhanceosome structure from^110^ shows mostly weak interactions that are likely to be transient, with only two pairs of TFs (highlighted in the grey box) having a buried surface area over 200 Å^2^. These weak interactions are of a similar magnitude to the 120 Å^2^ buried surface area between the two Tead4 proteins in the wild-type simulations (B, C, D, orange fill). **D**) Buried surface area distributions from four MD simulations of TEAD4 models. The values from the high-affinity double motif are shown as solid areas, and the left and right motif mutations are in shades of red. Upon disruption of the left component of the *Tead double* motif, the protein-protein interactions are weakened. Interestingly, upon mutating the right component, the left protein dissociates from the nucleobases in its motif entirely and forms a stable dimer with the remaining TEAD4 molecule. These two mechanisms point to the same conclusion: Weakening either motif disrupts the whole complex. **E**) An MD simulation with a low-affinity double motif of putative *Amotl2* enhancer tells a similar story to the mutated high-affinity motif (Figure S4D). The areas of measured interactions between protein-DNA on two *Tead* components of the low-affinity double motif are shown. The protein-nucleobase interactions are weakened (red line), and the protein-protein interaction becomes slightly weaker (red line), suggesting that the whole complex is less stable than the high-affinity double motif (solid area). **F**) Buried surface area calculations from a high-affinity double motif (solid area) and single base insertion led to a less labile complex that did not show the large protein-protein interaction areas we saw during the wild-type simulation. **G**) BPNet predictions of TEAD4 footprint on the wild-type and either left or right mutated sequences of the high-affinity double motif from putative *Tjp1* enhancer. Mutated nucleotides for left or right sequences are highlighted and were chosen based on the lowest contribution score at the position. **H**) BPNet predictions of TEAD4 footprint on the wild-type and either left or right mutated sequences of the low-affinity double motif from putative *Amotl2* enhancer. Mutated nucleotides for left or right sequences are highlighted and were chosen based on the lowest contribution score at the position. **I**) The contribution score sum was calculated at the low-affinity *Tead double* motif and on either side of the *Tead* component of the double motif. **J**) Luciferase assay of the wild type and mutated 175 bp minimal putative enhancer of *Amotl2* (mm10- *chr9:102,726,395-102,726,570*) were performed in three biological replicates and normalized to the empty vector control. Significance was determined by a student’s t-test (*p* < 0.05). **K**) The metapeak of observed and predicted TEAD4 binding at the mapped *Tead double* motifs shows BPNet learned *double* motifs in two cell types. *Tead double* motifs (TSCs:- ∼14k and ESCs:- ∼1k) were from fold 1 of their respective trained models. **L**) The *Tead double* motifs were discovered with TF-MoDISco with additional models trained with the same architecture as part of three-fold validation (fold 2 and fold 3). Spearman counts correlation values (top right) were determined by comparing the observed ChIP-nexus counts with BPNet’s predicted counts at TEAD4 ChIP-nexus peaks in ESCs.

**Figure S5.**
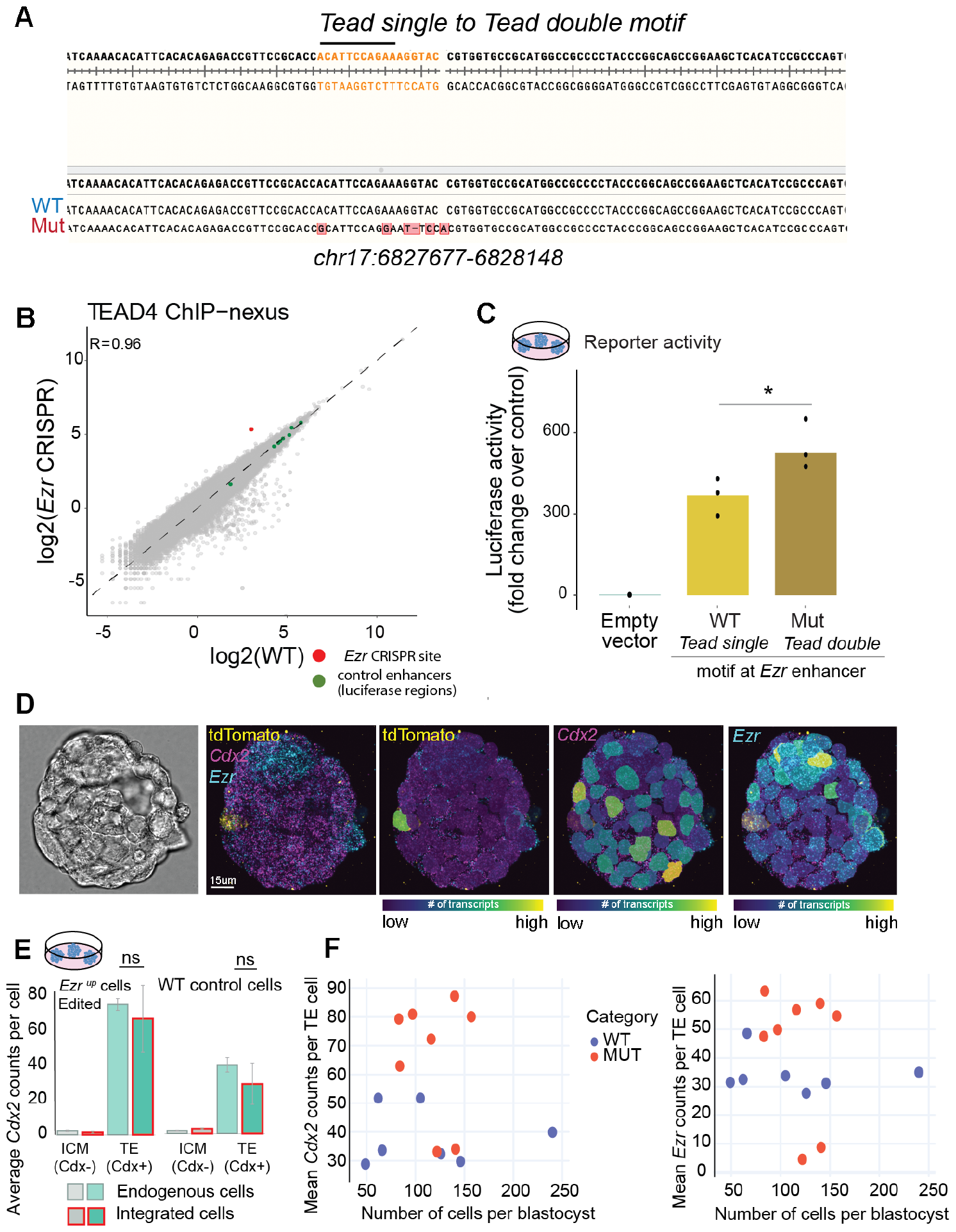
The edited *Tead double* motif within *the Ezr* enhancer shows increased TEAD4 binding, activity, and cell-specific gene expression (related to Figure 6) **A**) Sanger sequencing chromatogram at the *Ezr* enhancer with wild-type (WT) *Tead single* motif replaced/edited with the *Tead double* motif via homology-directed CRISPR-Cas9. **B**) Pairwise comparisons for TEAD4 ChIP-nexus between WT and CRISPR clone cells show high Pearson correlations for control enhancers (green dots) from the luciferase assay (Figure S2B), and the selected *Ezr* enhancer (CRISPR site in red dot) were consistent between the two replicates. **C**) Luciferase assay of the wild type and mutated 200 bp minimal *Ezr* enhancer were performed in three biological replicates and normalized to the empty vector control. Significance was determined by a student’s t-test (*p* < 0.05). **D**) HCR-FISH was performed on aggregated mouse blastocyst embryos with wild-type (WT) or mutant (edited Ezr^up^) cells for probes *Cdx2, Ezr*, and *td-Tomato* for quantification. The nuclei masks were made with Cellpose and Napari software using the DAPI channel, which was used on other channels to quantify average *Cdx2* counts to distinguish cells between inner cell mass and trophectoderm layer (shown in E), and td-Tomato stain was used to distinguish between native vs aggregated cells). **E**) Average quantification of *Cdx2* counts. Student’s t-test was performed between endogenous and integrated cells of edited Ezr^up^ and wild-type cell population (*p* > 0.05). **F**) The aggregated embryos with wild-type cells show an overall lower expression of average *Cdx2 or Ezr* expression with respect to embryo size (number of cells per blastocyst) than mutant (edited Ezr^up^) cells. All quantification was made per blastocyst to account for differences in expression.

